# Size evolution of gigantic genomes suggests stochastic outcomes of transposable element/host silencing interactions

**DOI:** 10.1101/2024.07.22.604708

**Authors:** Jie Wang, Guangpu Zhang, Cheng Sun, Liming Chang, Yingyong Wang, Xin Yang, Guiying Chen, Michael W. Itgen, Ava Haley, Jiaxing Tang, Rachel Lockridge Mueller

## Abstract

Size evolution among gigantic genomes involves gain and loss of many gigabases of transposable elements (TEs), sequences that parasitize host genomes. Animals suppress TEs using piRNA and KRAB-ZFP pathways. TEs and hosts coevolve in an arms race, where suppression strength reflects TE fitness costs. In enormous genomes, additional TE costs become miniscule. How, then, do TEs and host suppression invoke further addition of massive DNA amounts? We analyzed TE proliferation histories, deletion rates, and community diversities in six salamander genomes (21.3 - 49.9 Gb), alongside gonadal expression of TEs and suppression pathways. TE activity is higher in testes than ovaries, attributable to lower KRAB-ZFP suppression. Unexpectedly, genome size/expansion is uncorrelated with TE deletion rate, proliferation history, expression, and host suppression. Also, TE community diversity increases with genome size, contrasting theoretical predictions. TE/host antagonism in gigantic genomes likely produces stochastic TE accumulation, determined by noisy intermolecular interactions in huge genomes/cells.

## 1 Introduction

Transposable elements (TEs) are a diverse group of parasitic sequences that replicate and spread throughout host genomes using a variety of mechanisms, all of which include transcription (Wicker, et al. 2007; Lawlor and Ellison 2023). Found in all eukaryotes, their activity is suppressed by several host transcriptional and post-transcriptional silencing pathways (Czech and Hannon 2016; Deniz, et al. 2019; Almeida, et al. 2022). In animals, these include 1) the piRNA pathway, which comprises several small RNA-based gene silencing mechanisms, and 2) KRAB-ZFP-based silencing, in which KRAB-ZFP proteins bind specific TE sequences and recruit machinery to establish a transcriptionally repressive chromatin environment (Imbeault, et al. 2017; Ozata, et al. 2019). Both silencing pathways adapt to silence novel TE insertions. For TEs that do evade host silencing, each new insertion has several possible fates: if it confers a fitness advantage to the host, through regulatory or coding function, it can be a target of positive selection and become an integral part of the host genetic architecture. If it is strongly deleterious, it will be removed from the population by purifying selection. If it is neutral, or effectively neutral, its fate will be determined by genetic drift (Arkhipova 2018; Almeida, et al. 2022). TEs can be deleted from host genomes through large ectopic-recombination-mediated deletions or through smaller microdeletions (Bennetzen and Kellogg 1997; Petrov, et al. 2000).

Across eukaryotes, genomes vary both in TE abundance as well as the individual sequences that make up the genomic TE community; genomes can contain many or few TEs, and they can be diverse or homogeneous (Bourque, et al. 2018). In plants, the accumulation of huge numbers of TEs to achieve gigantic genome sizes is fairly common; there are thousands of plant genomes that exceed 10 Gb (Bureš, et al. 2024). In animals, on the other hand, genomic gigantism is rare, found in just a handful of clades. Within vertebrates, two clades have gigantic genomes – the lungfishes (6 species, genome sizes ranging from ∼40 to ∼130 Gb) and the salamanders (816 species, genome sizes ranging from ∼10 to ∼120 Gb) (AmphibiaWeb 2024; Gregory 2024). These two clades evolved gigantic genomes independently, and the two expansions appear to reflect different balances of evolutionary forces; lungfish genomes show signatures of increased strength of genetic drift, suggesting that ineffective purifying selection underlies the fixation of abundant TEs (Fuselli, et al. 2023). Salamanders, in contrast, do not show signatures of strong genetic drift (Mohlhenrich and Mueller 2016). Instead, they show evidence of low levels of deletion; both ectopic recombination-mediated deletion of large DNA segments, as well as small indels, are low in salamanders relative to genomes of more typical size, and this has been posited to explain (at least in part) the increase in genome size that they underwent ∼200 mya (Sun, et al. 2012; Sun and Mueller 2014; Frahry, et al. 2015; Liedtke, et al. 2018).

In gigantic genomes, the potential fitness consequences of any new individual TE insertion are far smaller than they are in compact genomes. The majority of sequence in gigantic genomes is non-essential, neither protein-coding nor regulatory (Novák, et al. 2020). Thus, novel TE insertions are much less likely to alter essential DNA sequences. In addition, any off-target spreading of epigenetic silencing of a novel TE insertion is less likely to affect essential sequences (Huang, et al. 2022), and given all the silenced TEs already in the genome, a novel TE insertion is likely to land in a transcriptionally repressed chromatin environment. In addition to the deleterious effects of insertion and silencing, TEs also harm host fitness by acting as non-homologous targets for recombination, yielding ectopic recombination events that can produce harmful deletions and duplications of essential sequences (Petrov, et al. 2011). Because gigantic genomes of salamanders experience low levels of ectopic recombination overall, their individual TEs have a lower chance of facilitating these harmful structural changes (Frahry, et al. 2015). Salamander genomes also have low rates of DNA loss through small indels as well as low rates of synonymous substitution, decreasing the chance of a neutral TE insertion mutating to a harmful gain-of-function allele (Sun, et al. 2012; Mohlhenrich and Mueller 2016). Taken together, these patterns demonstrate that the mutational hazard of individual TE insertions is low in gigantic salamander genomes. In the collective, the mass of DNA contributed by TE insertions has non-sequence-specific effects on organismal phenotype that are mediated through nucleus and cell size, which correlate positively with genome size (Gregory 2005; D’Ario, et al. 2021). Salamanders have low metabolic rates that are permissive to large cell sizes (Itgen, et al. 2022), and their cell biology and developmental systems are accommodating to large nucleus and cell volumes (Adams, et al. 2022; Taylor 2023; Taylor, et al. 2024). These cell-and organism-level properties of salamanders also lower the negative fitness consequences of TE insertions.

Host TE silencing pathways function to curb the harmful effects of TE proliferation, and hosts and TEs show patterns of sequence evolution consistent with coevolution between two entities that are in conflict, akin to an antagonistic host/parasite dynamic (Luo, et al. 2020; Zhang, et al. 2020; Lawlor and Ellison 2023). After a novel TE invades a genome, or alternatively after a TE evolves in sequence to escape host silencing, pressure is put on the host to curb that TE’s activity. In genomes of typical size, silencing of a new TE family will be favored by selection when the TE copy number reaches a level that harms host fitness enough to overcome genetic drift (Betancourt, et al. 2024). In gigantic genomes where each TE poses a lower risk, a much higher TE family size threshold has to be envisioned.

Across the salamander clade, genome size varies by an order of magnitude − from ∼10 Gb to ∼120 Gb (Gregory 2024). Even the smallest salamander genome is enormous compared to the vast majority of animal genomes. It is straightforward to imagine selection acting to silence a novel TE in a compact genome − and the arms race that would ensue as the TE evolves to evade this silencing − and to imagine that this might drive long-term trends in TE community and abundance. On the other hand, when considering genomes the size of salamanders’, it is less clear how any TEs could harm host fitness enough to elicit a selected response in host silencing. Thus, we hypothesize that the relationship between host silencing and TE abundance (and therefore genome size) might be different at the larger extremes of genome size than it is at the smaller extremes.

Here we present new data as well as detailed analyses of TEs and germline TE silencing in six species of diploid salamanders that differ by up to 28.6 Gb in haploid genome size, an amount equivalent to more than 9 human genomes. Using genomic sequence data, we summarize the diverse TE communities of each species as well as TE proliferation and deletion histories. Using RNA-seq data, we quantify TE expression in the gonads of both sexes as well as the expression of genes encoding proteins from two TE silencing pathways − piRNA and KRAB-ZFP. Using small RNA-seq data, we quantify the piRNA molecules that can guide associated protein complexes to silence TE activity in the gonads of both sexes. We integrate all three of these datasets to test whether the massive differences in TE-derived sequence abundance found among gigantic genomes reflect consistent differences in host TE silencing activity, TE deletion rates, TE community or insertion dynamics, or gonadal TE expression.

## 2 Results

### 2.1 Salamander genomes that differ by 28.6 Gb in size all contain numerous, largely overlapping active TE superfamilies

We generated and analyzed data from five species of salamanders whose genome sizes vary by 28.6 Gb (a 2-fold difference): four species from the family Salamandridae (*Tylototriton verrucosus*, genome size = 24.5 pg; *Pachytriton brevipes*, 39.8 pg; *Cynops orientalis*, 44.3 pg; and *Paramesotriton hongkongensis*, 51.0 pg) and one species from the family Cryptobranchidae (*Andrias davidianus*, 50.0 pg) (1 pg = 0.978 Gb; Fig. 1A) (Gregory 2024). We also incorporated previously published data for a species from the family Hynobiidae (*Ranodon sibiricus*, 21.8 pg) (Wang, et al. 2023) (Supplementary Table S1). This sampling spans the most basal split in the salamander phylogeny and flanks the median genome size (40 Gb) for the clade (Pyron and Wiens 2011; Gregory 2024). We performed low-coverage genome sequencing, yielding sequencing depths of 0.06X to 0.15X coverage (Supplementary Table S2). Following previous work, we used the PiRATE pipeline which was designed to mine and classify repeats from low-coverage genomic shotgun data in taxa that lack genomic resources (Berthelier, et al. 2018; Wang, et al. 2021; Wang, et al. 2023). The pipeline yielded 27,255*-*123,003 repeat contigs per species (Table 1). RepeatMasker mined the most repeats (18,449-102,254; 67.7-83.1%), followed by dnaPipeTE (2.4-11.9%), RepeatScout (4.4-8.6%), RepeatModeler (3.2-6.7%), or TE-HMMER (0-5.5%), depending on the species. Other methods found very few or no repeats (Table 1).

**Fig. 1.**
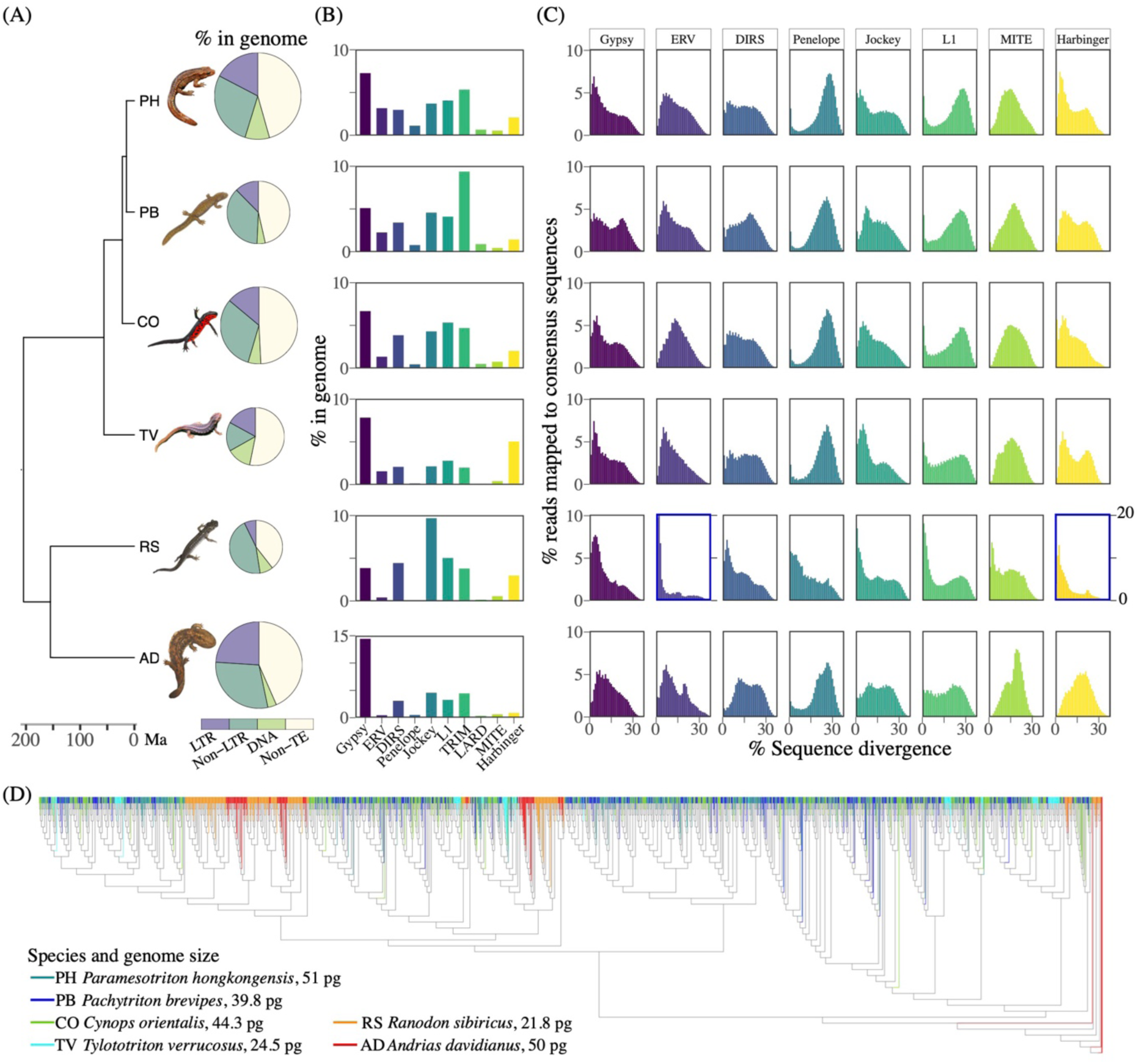
Genomic TE content, TE proliferation history, and phylogenetic relationships of an abundant TE superfamily in six salamander species (families Hynobiidae, Cryptobranchidae, and Salamandridae). (A) Phylogenetic relationships, divergence dates, genome sizes (reflected by the pie area), and genome content summarized at the level of TE order for the six taxa. The large proportions of non-TE sequences found in gigantic genomes result from the mutational decay of ancient TEs beyond recognition (Novák, et al. 2020). (B) Percentage of each genome made up of the 10 most abundant TE superfamilies or derivatives. (C) Proliferation histories of TE superfamilies represented by divergence between the TE sequences and the corresponding TE consensus sequence, with each bar representing 1% divergence. Note the different y-axis in the two panels highlighted in blue. (D) Unrooted phylogenetic tree showing the relationships among LTR/Gypsy genomic contigs; overall patterns are consistent with accumulation of this TE superfamily largely tracking the species tree.

**Table 1.** Count and percentage of the repeat contigs (≥ 100 bp, clustered at 100% identity) mined by different methods/software in the PiRATE pipeline (Berthelier, et al. 2018)

| TE-mining method | Software | *Tylotriton verrucosus* | *Pachytriton brevipes* | *Cynops orientalis* | *Andrias davidianus* | *Paramesotriton hongkongensis* |
| --- | --- | --- | --- | --- | --- | --- |
| Similarity-based | RepeatMasker | 18,449 (67.7%) | 92,791 (78.3%) | 102,254 (83.1%) | 47,417 (83.0%) | 88,948 (84.1%) |
|  | TE-HMMER | 1,317 (4.8%) | 4,248 (3.6%) | 6,782 (5.5%) | 0 | 4,509 (4.3%) |
| Structure-based | HelSearch | 1 (0.0%) | 433 (0.4%) | 2 (0.0%) | 169 (0.3%) | 3 (0.0%) |
|  | LTR harvest | 18 (0.1%) | 0 | 0 | 87 (0.2%) | 178 (0.2%) |
|  | MGEScan-non-LTR | 0 | 142 (0.1%) | 171 (0.1%) | 0 | 0 |
|  | MITE hunter | 3 (0.0%) | 9 (0.0%) | 12 (0.0%) | 3 (0.0%) | 8 (0.0%) |
|  | SINE finder | 9 (0.0%) | 56 (0.1%) | 70 (0.1%) | 22 (0.0%) | 101 (0.1%) |
| Repetitiveness-based | TEdenovo | 53 (0.2%) | 401 (0.3%) | 178 (0.1%) | 464 (0.8%) | 77 (0.1%) |
|  | RepeatScout | 2,336 (8.6%) | 5,394 (4.6%) | 6,675 (5.4%) | 4,134 (7.2%) | 4,643 (4.4%) |
| Repeat-building-based | dnaPipeTE | 3,243 (11.9%) | 11,061 (9.3%) | 2,945 (2.4%) | 2,087 (3.7%) | 3,482 (3.3%) |
|  | RepeatModeler | 1,826 (6.7%) | 4,000 (3.4%) | 3,914 (3.2%) | 2,696 (4.7%) | 3,770 (3.6%) |
| In total |  | 27,255 (100%) | 118,535 (100%) | 123,003 (100%) | 57,079 (100%) | 105,719 (100%) |
| Type of repeats annotated by PASTEC | Known TEs | 14,017 (51.4%) | 68,457 (57.8%) | 70,227 (57.1%) | 32,000 (56.1%) | 57,371 (54.3%) |
|  | Potential multi-copy host genes | 37 (0.1%) | 46 (0.0%) | 67 (0.1%) | 25 (0.0%) | 95 (0.1%) |
|  | Simple sequence repeats | 768 (2.8%) | 2,639 (2.2%) | 1,960 (1.6%) | 1,363 (2.4%) | 1,973 (1.9%) |
|  | Potential chimeras | 19 (0.1%) | 151 (0.1%) | 91 (0.1%) | 27 (0.0%) | 80 (0.1%) |
|  | Conflicting evidence | 1 (0.0%) | 5 (0.0%) | 7 (0.0%) | 6 (0.0%) | 0 (0.0%) |
|  | Unknown repeats | 12,413 (45.5%) | 47,237 (39.9%) | 50,651 (41.2%) | 23,658 (41.4%) | 46,200 (43.7%) |

Repeat contigs were annotated to TE order and superfamily in Wicker’s hierarchical classification system (Wicker, et al. 2007), modified to include several additional superfamilies, using PASTEC (Hoede, et al. 2014). Of the identified repeat contigs, 51.4% to 57.8% were classified as known TEs depending on species, representing 23 to 26 superfamilies in eight orders as well as retrotransposon and transposon derivatives (Supplementary Tables S3.1 to S3.6). <1% of contigs were filtered out as potential multi-copy host genes, chimeric contigs, or those with conflicting evidence. 1.6% -2.8% of contigs were filtered out as simple sequence repeats. Finally, 39.9% - 45.5% of repeat contigs were not classified (“unknown repeats” hereafter) (Table 1).

To calculate the proportion of different repeats in each species’ genome, shotgun reads were masked with RepeatMasker using two repeat libraries: excluding or including unknown repeats; in all cases, excluding unknown repeats increased the percentages of the genomes classified as known TE superfamilies (Supplemental Tables S3.1 to S3.6). For all species except *R. sibiricus*, LTR/Gypsy was the most abundant TE superfamily, comprising 7.83% (*P. brevipes*) to 23.37% (*A. davidianus*) of the shotgun read datasets (measured excluding unknown repeats). More generally, each of ∼10 TE superfamilies and/or TE derivatives contributed >1% of the data in each species, and there was much overlap in the identity of these “Top 10” despite some differences in rank abundance (Fig. 1C). Class I TEs are more abundant than Class II TEs in the genomes of all species, and LTR retrotransposons are more abundant than non-LTRs in all species except *R. sibiricus* (Fig. 1B; Supplementary Table S3.1-S3.6). The GC content of individual superfamilies ranged from 40% - 59% and the rank-order of GC content among superfamilies within each genome was broadly similar across species (Supplementary Figure S4). *P. hongkongensis* shows the strongest bias towards AT composition; GC content is < 50% in nine out of 10 TE superfamilies. In contrast, in the other species, four to six TE superfamilies have GC contents > 50%.

*R. sibiricus* showed patterns consistent with the highest ongoing levels of TE activity overall; five superfamilies are proliferating today at their highest-ever levels in its genome (ERV, Penelope, Jockey, L1, and MITE, demonstrated by the highest percentage of reads mapping to consensus sequences with < 1% sequence divergence) and no families appear inactive (Fig. 1C). In contrast, *A. davidianus* showed patterns consistent with the lowest ongoing levels of TE activity overall; three superfamilies appear inactive (DIRS, MITE, and Harbinger, demonstrated by < 1% of reads mapping to consensus sequences with < 1% sequence divergence), and no superfamilies are proliferating at their highest-ever levels. The four species of Salamandridae show evidence of similar TE activity overall; MITEs are inactive in all genomes, ERV is inactive or close to it in all genomes, and L1 activity is at its highest, or near-highest, today following a historic dip in activity. Although it is notable that the smallest genome (*R. sibiricus*) has the highest TE activity, and the near-largest genome (*A. davidianus*) has the lowest TE activity, similar patterns are not visible across the two-fold ranges of genome size within the Salamandridae clade (Fig. 1C).

### 2.2 Phylogeny of LTR/Gypsy TE superfamily sequences shows that TE diversification largely tracks the species tree

The phylogenetic relationships among LTR/Gypsy genomic contigs reveal clades that largely track the species tree; contigs from *R. sibiricus* and *A. davidianus* tend to cluster together, sometimes in reciprocally monophyletic clades (Fig. 1D), consistent with their membership in the suborder Cryptobranchoidea with a ∼151 mya divergence time (Fig. 1A). Similarly, sequences from the four species of Salamandridae tend to cluster together, with sequences from sister taxa *P. brevipes* and *P. hongkonesnsis* (divergence time ∼15 mya) frequently clustered together. We saw no strong signature of increased diversification of LTR/Gypsy contigs in *P. hongkongensis* following divergence from *P. brevipes*, despite a ∼10 Gb difference in genome size. The phylogeny was well-supported; 75.0% of nodes received bootstrap support values > 95% and 84.6% of nodes received bootstrap support values > 80%. Overall, our results suggest that the evolutionary accumulation of novel TE insertions, at least in the abundant LTR/Gypsy superfamily, largely tracks the species tree.

### 2.3 TE diversity increases with genome size in salamanders, unlike in other vertebrate groups

We selected 88 species of “fishes”, frogs, salamanders, non-avian reptiles, birds, and mammals and tested for the predicted negative relationship between TE diversity and genome size in each group (Wang, et al. 2021). Diversity of each genomic TE community was measured using both Gini-Simpson’s and Shannon diversity indices, considering TE superfamilies as “species” and the percentage of the genome occupied by each TE superfamily as “abundance”. The two diversity indices revealed similar patterns (Supplementary Table S5). Overall, the ectotherms had more diverse TE communities than the endotherms (Fig. 2). Neither birds nor mammals showed a relationship between genome size and TE community diversity. In contrast, 3 of the 4 ectothermic clades or groups showed a significant (frogs, salamanders) or near-significant (non-avian reptiles; *P* < 0.05 for Shannon and = 0.07 for Gini-Simpson) relationship between genome size and TE diversity. In the frogs and non-avian reptiles, TE diversity is higher in smaller genomes, as predicted by several theoretical models (Wang, et al. 2021). However, salamanders showed the opposite pattern; TE diversity increases with increasing genome size, in contrast with theoretical predictions. The other species with gigantic genomes − the lungfishes − had comparable TE diversity levels with the smallest salamander genomes. There was no phylogenetic pattern to the TE diversity levels across salamanders; family-level clades had overlapping ranges of diversity indices (Supplementary Table S5). Consistent with these broader results, within our six focal salamander species, TE diversity increases monotonically with genome size with the exception of *A. davidianus*, which has diversity levels comparable to other salamanders with smaller genome sizes.

**Fig. 2.**
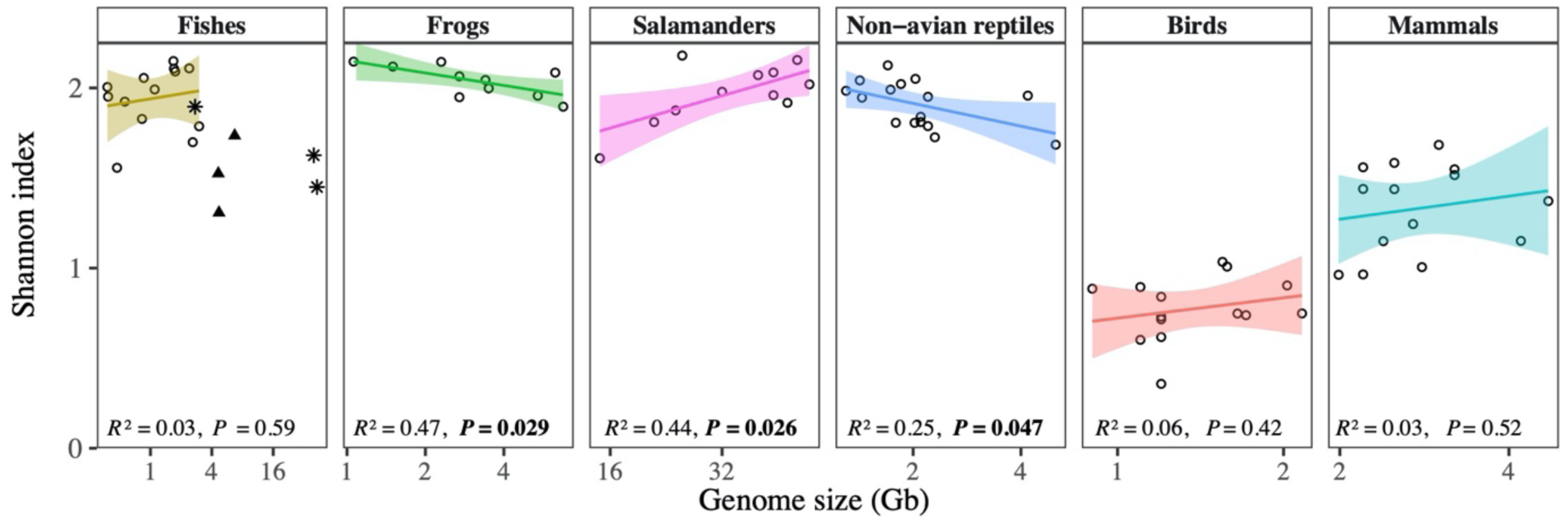
Shannon diversity indices summarizing the TE communities from vertebrate genomes. Shaded regions represent the 95% confidence intervals. In the “fishes”, 〇 are ray-finned fishes, ▴ are cartilaginous fishes, and ✱ are non-tetrapod lobe-finned fishes (coelacanth and lungfishes); the regression line is just for the ray-finned fishes. Salamanders show a significant positive relationship between genome size and diversity, in contrast to other groups.

### 2.4 Large deletions of LTR/Gypsy sequences are equally prevalent across salamander genome sizes

LTR retrotransposon structure includes terminal repeats that can undergo ectopic recombination, eliminating the internal sequence and one copy of the terminal repeat sequence. These deletions produce diagnostic solo LTR sequences, resulting in elevated abundances of terminal sequences relative to internal sequences genome-wide; higher ratios of terminal to internal sequences indicate higher levels of deletion. Within each salamander species, the ratio of terminal to internal sequences (T:I) across individual LTR/Gypsy element families varied roughly 10- to 30-fold (Fig. 3); the lowest ratio of any LTR/Gypsy family was found within *C. orientalis* (T:I = 0.15), whereas the highest was found in *A. davidianus* (T:I = 8.6). However, across species, there was no significant difference in the T:I ratio of LTR/Gypsy elements (ANOVA, *F*_4,85_ = 2.11, *P* = 0.087) (Fig. 3). The species medians (0.43 – 0.94) were comparable to previously published estimates for salamanders of the family Plethodontidae (0.55 – 1.25), but much lower than those in the genomes of other non-salamander vertebrates (Frahry, et al. 2015). Thus, we see no evidence that deletion rates of this abundant TE superfamily are lower in larger salamander genomes than in smaller salamander genomes.

**Fig. 3.**
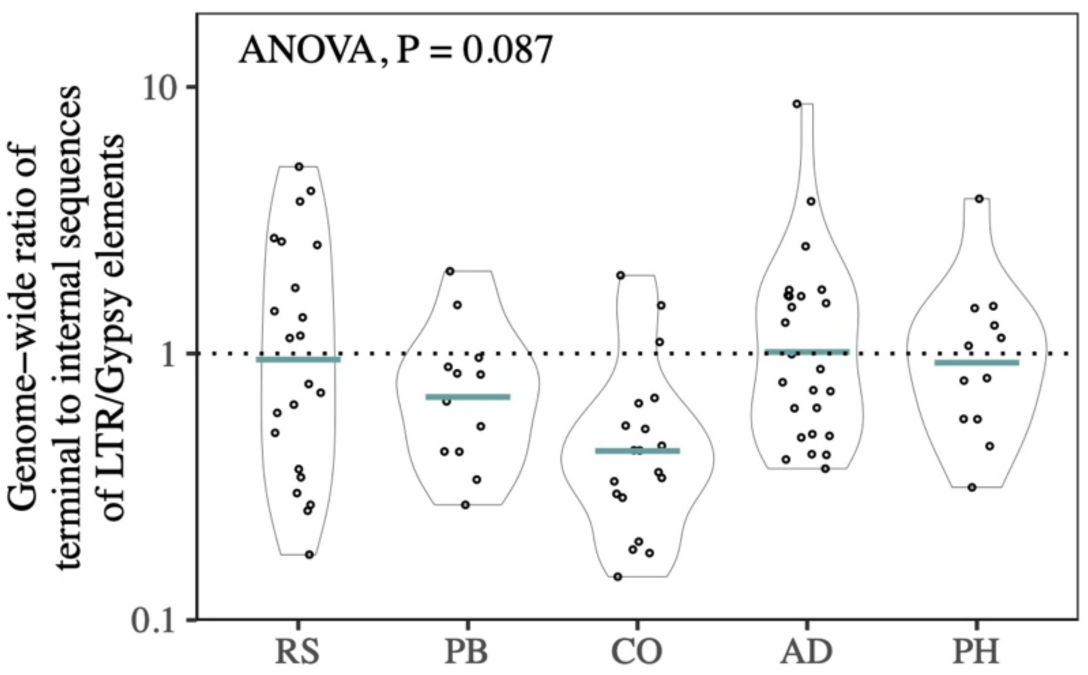
Ectopic recombination-mediated deletion levels of LTR retrotransposons are the same across the genomes of salamanders. Deletion levels are measured as the genome-wide ratio of terminal to internal sequences of LTR/Gypsy elements. A ratio of 1:1 (dotted line) is expected in the absence of ectopic recombination-mediated deletion of LTR/Gypsy sequences from the genome. Species are arranged left to right by increasing genome size: RS, *Ranodon sibiricus*; PB, *Pachytriton brevipes*; CO, *Cynops orientalis*; AD, *Andrias davidianus*; PH, *Paramesotriton hongkongensis*. The y-axis is on the log scale.

### 2.5 Gonadal TE expression is correlated with genomic abundance in all species and is consistently higher in testes than ovaries

In all focal salamander species except *R. sibiricus*, TE expression levels for both males and females were significantly positively correlated with genomic abundance at the TE superfamily level based on linear regression (*P* ≤ 0.016; Fig. 4A). Across species, the slopes of the linear regressions were not significantly different from one another in either sex (*P* > 0.59), but the intercepts were different in both sexes (*P* < 0.001) (Fig. 4E; Supplementary Table S6). Slopes were not significantly different between males and females in any species (*P* > 0.5), but intercepts were higher in males than females in *T. verrocosus* and *A. davidianus* (*P* < 0.04) (Fig. 4A; Supplementary Table S6).

**Fig. 4.**
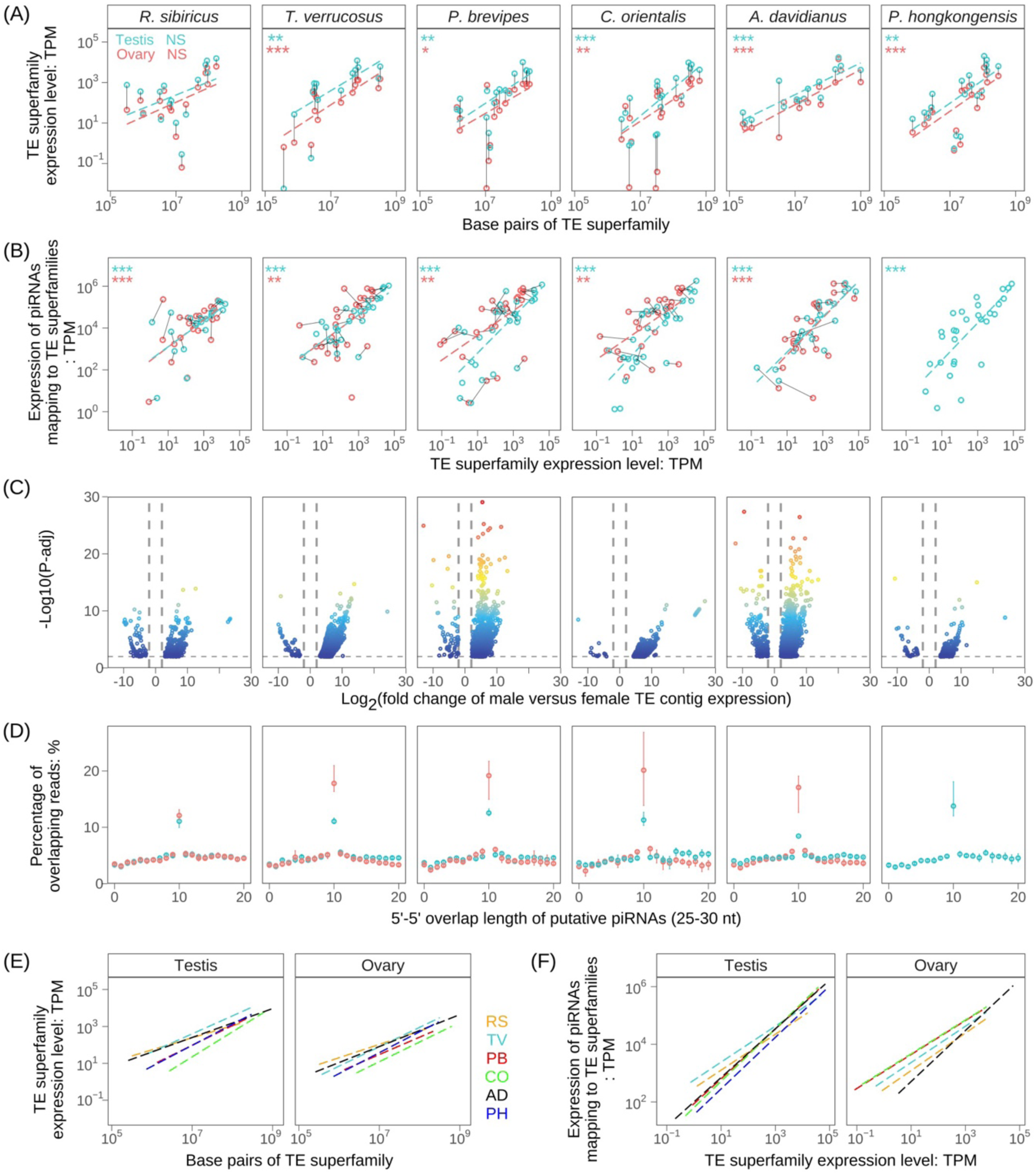
(A) Positive correlations between abundance and gonadal expression for TE superfamilies. In all species except *R. sibiricus*, expression and genomic abundance are significantly correlated. Gray lines connect the male and female expression values for each TE superfamily; male expression is higher than female expression for the majority of superfamilies. NS means not significant; *, **, and *** mean *P* < 0.05, < 0.01, and < 0.001, respectively. (B) Positive correlations between TE superfamily expression and mapping piRNAs in testes and ovaries. (C) Volcano plots showing differential expression between males and females for individual TE contigs; positive fold change values indicate male-biased expression, and negative fold change values indicate female-biased expression. Four contigs from *A. davidianus* and two contigs from *P. brevipes* with extreme adjusted *P*-values were omitted for visual clarity. (D) The percentage of 5’ -5’ overlap between pairs of sense and antisense piRNAs of different lengths in testes and ovaries. The high Z-score associated with the 10 nt overlap is a typical signature of ping-pong piRNA biogenesis. (E) Linear regression lines showing the correlations between abundance and gonadal expression for TE superfamilies for all six species plotted together for testes (left) and ovaries (right). (F) Linear regression lines showing the correlations between TE superfamily expression and the expression of piRNAs mapping to TE superfamilies of all six species plotted together for testes (left) and ovaries (right). For (E) and (F), slopes are not significantly different among species for either sex, but intercepts are significantly different for both sexes (Supplementary Table S6, S12).

Expression was higher in the male gonads than the female gonads for the majority of TE superfamilies in all species (Fig. 4A). In three species (*A. davidianus*, *P. hongkongensis,* and *R. sibiricus*), males and females shared the most highly expressed TE superfamily (DIRS, DIRS, and Jockey, respectively) (Supplementary Table S7). In the remaining three species, males and females had a different most highly expressed TE superfamily in the gonads (either DIRS/DIRS, LINE/L1, or LTR/Gypsy). *R. sibiricus* was the only species for which the most abundant TE superfamily was also the mostly highly expressed in both males and females; in all other species, the most highly expressed TE superfamily is among the top five most abundant in the genome. We identified varying numbers of differentially expressed TE contigs between male and female gonads across species, with some species showing a greater skew in the number of contigs biased towards male versus female expression: *R. sibiricus*, 463 contigs (360 male-biased, 103 female-biased); *T. verrucosus*, 2,703 contigs (2,661 male-biased, 42 female-biased); *P. brevipes*, 2,668 contigs (2,606 male-biased, 62 female-biased); *C. orientalis*, 1,167 contigs (1,154 male-biased, 13 female-biased); *A. davidianus*, 2,918 contigs (2,670 male-biased, 248 female-biased); and *P. hongkongensis*, 321 contigs (286 male-biased, 35 female-biased) (Fig. 4C). For both sexes in all species, transcripts annotated as endogenous protein-coding genes were at least an order of magnitude more highly expressed in the gonadal transcriptome than TE transcripts, but TEs comprised a 1.7 – 4.2- fold greater percentage of the transcriptome in males than in females (Supplementary Figure S8). Contigs that received no gene or TE annotation (i.e. those associated with pervasive transcription) were also more highly expressed in males than in females. Percentages of gonadal transcriptomes composed of TEs were comparable between these salamanders and other vertebrates (Pasquesi, et al. 2020). Our results reveal no consistent differences in gonadal expression of TE superfamilies that correlate with increasing genome size across our focal salamanders.

### 2.6 The pool of unique piRNAs that map to TEs has similar overall characteristics in testes and ovaries and across species, but individual sequences are different

For all species, in both testes and ovaries, the length distribution of small RNA molecules includes a peak at 29 or 30 nucleotides (Supplementary Figure S9A), and sequences with lengths of 25-30 nt show a strong 5’-U bias at the first nucleotide position, consistent with expectations for the piRNA pool (Supplementary Figure S9B). A second peak at 22 nt also appeared in some species (in ovaries: *Ranodon*; in testes: *Cynops*, *Paramesotriton*, and *Andrias*, where it was higher than the peak at 29 or 30 nt). Molecules with lengths of 25-30 nt were treated as putative piRNAs and subjected to additional analyses; molecules in this size range comprise 37% (*R. sibiricus*) to 53% (*T. verrucosus*) of the total 18-40 nt reads in females and 18% (*A. davidianus*) to 60% (*R. sibiricus*) of the total 18-40 nt reads in males (Supplementary Table S10).

The average normalized number of unique piRNA sequences in each individual ranges from 460,533 (*T. verrucosus*) to 663,310 (*A. davidianus*) for females and from 366,003 (*A. davidianus*) to 708,587 (*P. brevipes*) for males. Female and male numbers of normalized unique piRNAs overlap in all species except *P. brevipes*, where the highest female value is ∼2% lower than the lowest male value (Supplementary Table S10).

The average percentage of unique piRNAs that map to TE transcripts in each individual ranges from 14.2% (*P. brevipes*) to 29.6% (*T. verrucosus*) for females and from 16.1% (*P. brevipes*) to 30.2% (*T. verrucosus*) for males (Table 2). Female and male numbers of normalized unique piRNAs that map to TEs overlap in all species except *P. brevipes* (lowest male ∼10% greater than highest female) and *A. davidianus* (lowest female ∼6% greater than highest male) (Supplementary Table S10). Across species, numbers of unique piRNAs that map to TEs broadly overlap and show no trend with increasing genome size. Although all six genomes contain largely overlapping “Top 10” TE superfamilies, including some similarities in rank abundance, the individual sequences in the pool of unique piRNAs show very little overlap across species (Supplementary Figure S11). These patterns are consistent with a broadly stable community of TE lineages and silencing pathways that show evidence of constant evolutionary change, consistent with coevolution between two antagonistic entities.

**Table 2.** Unique piRNA sequences, and percentages mapping to TEs, in male and female gonads.

| Species | Normalized average number for testes | Average percentage mapping to TEs for testes | Normalized average number for ovaries | Average percentage mapping to TEs for ovaries |
| --- | --- | --- | --- | --- |
| *R. sibiricus* | 490,553 | 20.73% | 462,804 | 22.75% |
| *T. verrucosus* | 664,382 | 30.24% | 460,533 | 29.55% |
| *P. brevipes* | 708,587 | 16.12% | 585,166 | 14.16% |
| *C. orientalis* | 572,251 | 16.97% | 624,442 | 16.16% |
| *A. davidianus* | 366,003 | 20.12% | 663,310 | 25.52% |
| *P. hongkongensis* | 554,138 | 20.13% | — | -- |

### 2.7 TE-mapping piRNA expression is correlated with TE expression in ovaries and testes in all species

In both testes and ovaries of all species, TE expression at the superfamily level is significantly positively correlated with the expression of mapping piRNAs based on linear regression (*P* ý 0.008) (Fig. 4B). Across species, the slopes of the linear regressions were not significantly different from one another in either sex (*P* ≥ 0.27), but the intercepts were different in both sexes (*P* < 0.001) (Fig. 4F; Supplementary Table S12). The slopes of the linear regressions were not significantly different between the gonads of the two sexes in any species, but the intercept in males was higher than females in *C. orientalis* (Supplementary Table S12). In both sexes across all species, only half or fewer of the expressed TE contigs are mapped by any piRNAs (Table 3). Our results reveal no consistent differences in gonadal expression of TE-mapping piRNAs relative to TE expression that correlate with increasing genome size across our focal salamanders.

**Table 3.** Expressed TE contigs mapped by piRNAs in male and female gonads.

| Species | Average number<br>for testes | Average percentage mapped<br>by piRNAs for testes | Average number<br>for ovaries | Average percentage mapped<br>by piRNAs for ovaries |
| --- | --- | --- | --- | --- |
| *R. sibiricus* | 12,812 | 28.7% | 17,111 | 38.3% |
| *T. verrucosus* | 17,717 | 49.6% | 13,514 | 37.8% |
| *P. brevipes* | 12,704 | 50.5% | 12,163 | 48.3% |
| *C. orientalis* | 14,043 | 31.3% | 13,100 | 29.2% |
| *A. davidianus* | 10,233 | 45.1% | 14,124 | 45.1% |
| *P. hongkongensis* | 19,054 | 30.7% | — | -- |

### 2.8 Ping-pong amplification of TE-mapping piRNAs is more pronounced in ovaries than testes

In all five species for which we have both male and female piRNA data, the piRNA pool includes the signature of ping-pong amplification, and the signature is more pronounced in the female gonads than in the male gonads (Fig. 3D). Overall, there is no clear trend associating ping-pong amplification with increasing genome size in either males or females.

In all six species, we identified varying numbers of putative piRNA cluster transcripts in both males and females (8 - 22 putative transcripts in males, 9 – 22 putative transcripts in females, contig lengths 2,165 – 15,954 nt). In all species except *C. orientalis*, the majority were annotated with at least one TE insertion and a minority (or none) were annotated to contain an endogenous protein-coding gene. Testes showed higher and more variable expression than ovaries for all species, which would suggest higher overall piRNA cluster transcript expression in male gonads (Supplemental Figure S13). However, given the small percentage of putative piRNAs that were found within one of these putative piRNA cluster transcripts (0.7% - 2.9% depending on species), as well as the small fraction of the overall genome size encompassed by the length of these putative transcripts (<0.0008%), we adopt a conservative approach to these results and do not interpret them further. We encourage further exploration of piRNA cluster transcripts in gigantic genomes.

### 2.9 Expression of TE silencing pathway genes differs between testes and ovaries and across species

piRNA pathway gene expression (relative to miRNA pathway gene expression) is significantly different between males and females in all species except *C. orientalis* (*P* < 0.018) and is significantly different across species for both sexes (ANOVA, *P* ≤ 0.003). In three species, piRNA pathway gene expression is higher in testes than in ovaries (Fig. 5A), whereas for two − *A. davidianus* and *P. hongkongensis*, which have the largest genome sizes expression is higher in ovaries than in testes. In males, the highest piRNA pathway expression is found in *R. sibiricus*, which has the smallest genome size as well as the highest overall TE activity based on proliferation histories. The lowest male piRNA pathway expression levels are found in *A. davidianus*, which has the lowest overall TE activity based on proliferation histories and close to the largest genome size, and in *P. hongkongensis*, which has the largest genome size (Fig. 1C). The remaining three species (all from the family Salamandridae) have intermediate male piRNA pathway expression. Female piRNA pathway expression (relative to miRNA pathway gene expression) shows no relationship with overall TE activity or genome size, but it is notable that the species with the lowest male piRNA pathway expression (*A. davidianus*) has the highest female piRNA pathway expression.

**Fig. 5.**
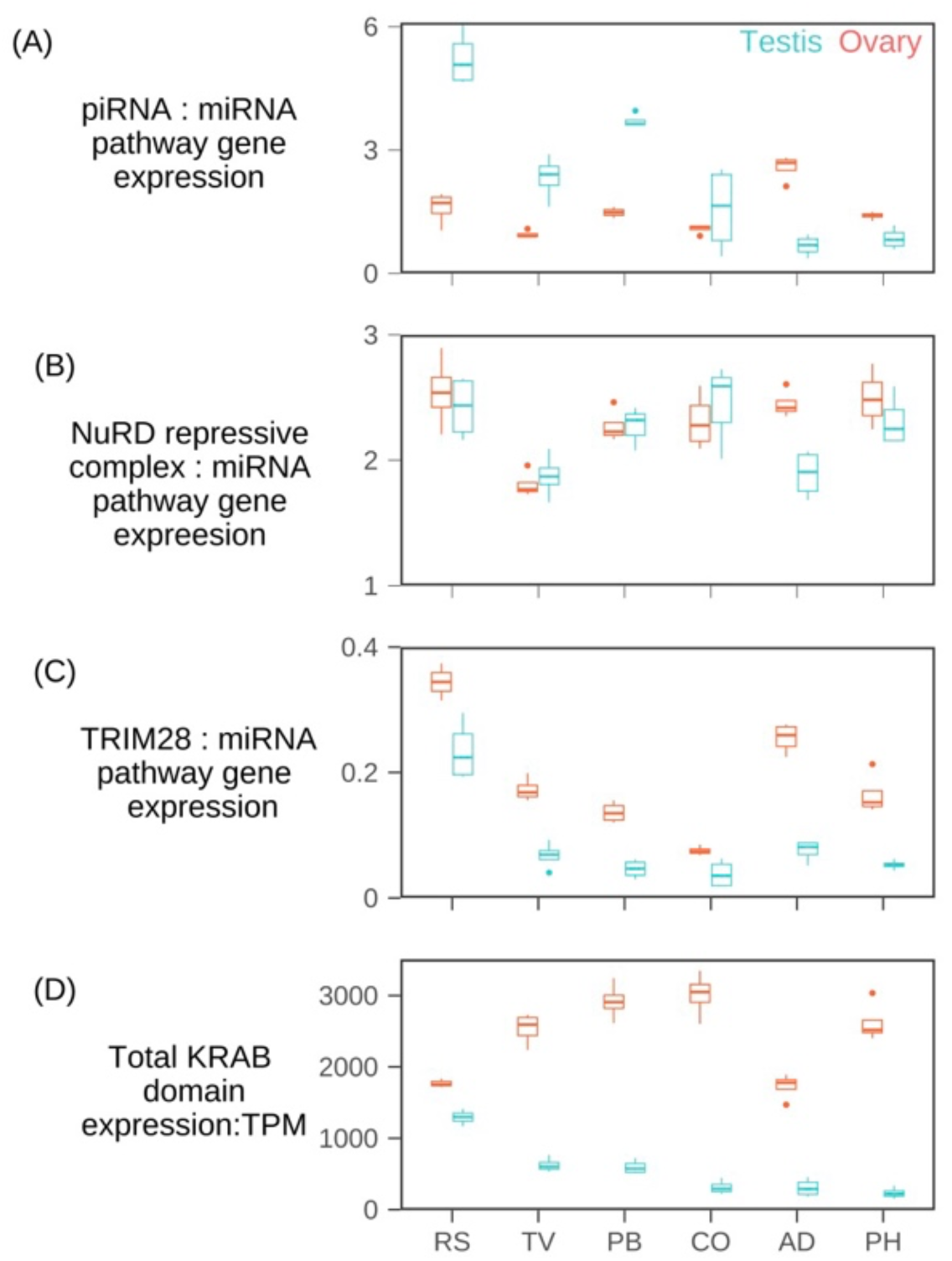
Ratio of the summed expression of (A) piRNA pathway genes, (B) NuRD and associated repressive complex genes, and (C) TRIM28 to the summed expression of miRNA pathway genes across salamanders. Note the different y-axes. (D) Summed expression of all KRAB domains. Species are arranged left to right by increasing genome size: RS, *Ranodon sibiricus*; TV, *Tylototriton verrucosus*; PB, *Pachytriton brevipes*; CO, *Cynops orientalis*; AD, *Andrias davidianus*; PH, *Paramesotriton hongkongensis*.

NuRD repressive complex gene expression (relative to miRNA pathway genes) is similar between males and females in five of the six species; the exception is *A. davidianus*, where female expression is significantly higher than male expression (*P* < 0.001). Across species, expression is significantly different for both sexes (*P*:: 0.009), but with no clear relationship with genome size or overall TE activity.

TRIM28 expression (relative to miRNA pathway expression) is significantly higher in ovaries than in testes for all six species (*P* < 0.038) and is significantly different across species for both sexes (ANOVA, *P* < 0.001). Male and female expression is highest in *R. sibiricus* (with its high overall TE activity and low genome size), while the remaining five species show no relationship with genome size or overall TE activity. KRAB domain expression is also significantly higher in ovaries than in testes for all six species (*P* < 0.001) and is significantly different across species for both sexes (ANOVA, *P* < 0.001). KRAB domain expression in testes declines with increasing genome size, whereas no clear relationship exists between KRAB expression and genome size or overall TE activity in ovaries.

## 3 Discussion

*TE expression is higher in males in all species, likely reflecting a sex difference in KRAB-ZFP-mediated TE silencing rather than piRNA-mediated TE silencing*.−Across all six focal salamander species, TE expression is higher in the male gonads than in the female gonads − testes have more TE transcripts per genomic abundance than ovaries for the majority of TE superfamilies, more TE contigs show testis-biased expression than ovary-biased expression, and the overall percentage of the transcriptome made up of TEs is higher in testes than in ovaries. This pattern of male-biased TE expression suggests that TE silencing is less effective in the male germline, and patterns consistent with this prediction (i.e. lower TE-mapping piRNA expression, lower piRNA pathway gene expression, and weaker ping-pong signature in males) have been interpreted in this way in *Drosophila* (Saint-Leandre, et al. 2020). In our focal salamanders, however, the situation is less straightforward. Females have higher expression of TRIM28 and KRAB domains in all six species, consistent with stronger KRAB-ZFP-mediated TE silencing in ovaries than in testes. However, there is no consistent difference between females and males in many piRNA-related metrics of TE silencing: on the one hand, females do have a stronger signature of ping-pong amplification than males. On the other hand, the piRNA pool itself (e.g. the number of unique piRNA sequences mapping to TEs) and the extent to which the piRNA pool targets TEs (e.g. the number of TE transcriptome contigs that are mapped by piRNAs, as well as the mapping density of piRNAs to TE transcripts) are either not different between males and females, or are different in only a subset of species with males or female showing the higher value. In addition, piRNA pathway genes are more highly expressed in males in four of the six species and more highly expressed in females in the remaining two, and NuRD repressive complex genes are expressed at equal levels in five of the six species. In sum, our data reveal consistent differences between the sexes in TE expression in salamander gonads (in contrast to (Carducci, et al. 2021)). The molecular mechanisms underlying this difference require further study, and we suggest based on our TRIM28 and KRAB domain expression results that these mechanisms include higher activity of KRAB-ZFP-mediated silencing in ovaries. We note, however, that levels of transcription of unannotated contigs (i.e. pervasive transcription) are also higher in male gonads than in female gonads (Supplementary Figure S7), which could be indicative of a more generally accessible chromatin state in testes that would allow higher TE transcription.

*Deletion, diversity, and GC content suggest that salamander TE silencing pathways do not strongly antagonize the TE community.−* Ectopic recombination is one of the molecular mechanisms that deletes TEs from individual genomes and, thus, contributes low-TE variants to a population, allowing genomes to evolve to smaller sizes (Yu, et al. 2024). On the other hand, ectopic recombination is also one of the primary negative fitness consequences of TE loci because off-target recombination can produce harmful deletions and duplications of intervening functionally important coding sequences (Petrov, et al. 2003; Furano, et al. 2004). Therefore, genomes with lower propensity towards ectopic recombination have been hypothesized to be more tolerant of TEs, leading to increased TE accumulation and genome size (Furano, et al. 2004). Several models have proposed that, as TEs accumulate and genomes expand, the diversity of the overall TE community should decrease for different (but not necessarily mutually exclusive) reasons (Wang, et al. 2021): Petrov (2003) posits that selection for diverse TE communities is simply relaxed in larger genomes because of their lower rates of ectopic recombination. Furano (2004) posits that TEs can accumulate and be more active in genomes with lower rates of ectopic recombination, which leads to out-competition of many TE lineages by the lineage that most successfully exploits the host’s replication factors. Boissinot and Sookdeo (2016) posits that the higher TE abundance and activity tolerated by genomes with lower ectopic recombination trigger an arms race between TEs and hosts, which leads to a decrease in diversity (i.e., host silencing results in only one TE family active at a time) (Boissinot and Sookdeo 2016). Finally, the models used in Abrusán et al. (2006) suggest that increased genome size with more active TE copies would lead to lower TE diversity if there were non-sequence-specific cross-reactive silencing among TE families, which would lead to competition to evade silencing among the TEs and thus decreased diversity (Abrusán and Krambeck 2006).

Salamanders have low rates of ectopic recombination, one of the explanations for TE accumulation and genome expansion in the clade (Frahry, et al. 2015). In contrast to all of these predictions, however, salamanders show increased rather than decreased TE diversity at the superfamily level as their TE abundance and genome size increase (Figure 2), and diversity at the TE family level remains high across genome sizes (Supplementary Files 3.1-3.6; see the number of genomic contigs in each TE superfamily after CD-hit clustering at 80% similarity, a proxy for Wicker’s 80/80/80 TE nomenclature rule (Wicker, et al. 2007; Wang, et al. 2021). These patterns suggest that 1) TEs are not in strong competition for access to rate-limiting host replication factors, 2) the arms race between TE proliferation and host silencing does not restrict activity to a single TE family at a time, and/or 3) TEs are not strongly competing to evade non-sequence-specific cross-reactive silencing mechanisms in salamanders. All of these scenarios are consistent with a genomic environment in which TE silencing machinery is not putting such substantial constraints on TE activity that it drives a strong coevolved response at the TE community level, despite TEs and hosts engaging in a coevolutionary arms race. Similarly, TE superfamilies in salamanders do not show the low GC content and high AT content characteristic of many TEs (Supplementary File S4). High AT content has been interpreted in other genomes to reflect both the TEs evading CpG methylation-based silencing as well as the TEs showing evolved “self-regulation” towards lower transcriptional efficiency (and therefore lower deleterious effects on the host) because high AT content is associated with slower transcription, RNA polymerase stalling and dissociation, and premature transcript termination (Boissinot 2022). Thus, the GC content patterns in salamanders, like the increasing diversity with increasing genome sizes, also point to a genomic environment in which host silencing is not driving a strong evolved response at the TE community level.

*How do TE activity and permissive TE silencing interact to shape genome expansion in gigantic genomes?−* Deletion rate can drive genome size evolution, but we see no evidence that it differs across these salamander genome sizes. Variation in the strength of genetic drift can also drive genome size evolution, but salamanders do not show patterns consistent with drift-mediated genome expansion (Rios-Carlos, et al. 2024). Thus, we focus on TE proliferation. Across the ∼30 Gb range of genome sizes encompassed by our focal taxa, we see no consistent, monotonic relationships between genome size and any measure of TE activity or host TE silencing; the pattern that comes closest is the expression of piRNA pathway genes in the testes, which is highest in the smallest genome and lowest in the largest genomes, but this pattern is not mirrored in the ovaries. However, TE activity and host silencing activity have also been hypothesized to differ based on whether the host or the TE community is currently “on the attack” in the TE/host arms race (Wang, et al. 2023). A reasonable proxy for a salamander lineage’s position along this oscillating evolutionary trajectory is whether genome size is shrinking, expanding, or remaining roughly constant. As an approximation for whether our focal taxa are likely experiencing a long-term trend towards genome expansion or contraction, we examined the known genome sizes for other members of the same families.

Four of our focal taxa are members of the family Salamandridae, which includes 139 species in 21 genera (AmphibiaWeb 2024). Genome size estimates for 45 species across 16 genera range from 21 to 51 pg. Three of our focal taxa − *P. hongkongensis* (51 pg), *P. brevipes* (40 pg), and *C. orientalis* (44 pg) *−* are nested within a small clade with the largest measured genome sizes within Salamandridae (≥ 40 pg), suggesting that all three lineages have been expanding (to varying degrees) since their divergence from the remainder of Salamandridae:: 50 mya (Kieren, et al. 2018; Gregory 2024). Thus, we postulate that these three gigantic genomes may represent lineages where the TE community is “on the attack.”

In contrast, *T. verrucosus* (24.5 pg) is nested within a small clade with some of the smaller genome sizes within Salamandridae (21-27 pg), and its phylogenetic position is consistent with either 1) maintenance of a small genome size (if the 21 pg genome of basal *Salamandrina terdigitata* reflects a small ancestral genome size for the family) or 2) being part of a smaller clade undergoing genome contraction since their divergence from the remainder of Salamandridae:: 60 mya (Kieren, et al. 2018; Gregory 2024). Similarly, *R. sibiricus* is a member of Hynobiidae, which includes 98 species in 9 genera. Genome size estimates for 9 species across 5 genera (*Onychodactylus fischeri*, *Salamandrella keyserlingi*, *Batrachuperus mustersi*, and five species of *Hynobius*, in addition to *R. sibiricus*) range from 16 to 25 pg except for *O. fischeri* (47.5 pg) (Pyron and Wiens 2011; AmphibiaWeb 2024; Gregory 2024), suggesting that *R. sibiricus* (21.8 pg) is likely not experiencing a strong directional trend in genome size. Finally, *A. davidianus* is a member of Cryptobranchidae, which includes 6 species in two genera. Genome size estimates for three species (*Cryptobranchus alleganiensis, Andrias japonicus,* and *A. davidianus*) range from 46.5 – 55 pg; thus, *A. davidianus* (50 pg) is also likely not experiencing a strong directional trend in genome size (Pyron and Wiens 2011; AmphibiaWeb 2024; Gregory 2024).

In the three focal genomes where TEs are most likely to be “on the attack” based on increasing genome size (*P. hongkongensis*, *P. brevipes*, and *C. orientalis*), we might logically predict higher levels of TE activity. In contrast, based on our plots of TE divergence (Figure 1C), *Ranodon sibiricus* shows the pattern most consistent with the highest TE activity, despite its likely static genome size − all TE superfamilies are active, and activity is at a historic high in five superfamilies. Based on our analysis of TE superfamily expression per genomic abundance, our three likely “TE-attacking” genomes actually exhibit the lowest TE expression/TE bp in both testes and ovaries (Figure 4E). Finally, our three likely “TE-attacking” genomes do not have the highest percentage of their gonadal transcriptomes made up of TEs. We note that the predictions for TE expression levels are complex, however; models suggest that, with additional TE insertions and genome expansion, a smaller proportion of genomic TE insertions will be active (Kijima and Innan 2013; Roessler, et al. 2018). Furthermore, TE transcript expression is a reasonable proxy for TE activity, but it is only one step of the TE life cycle; it is necessary, but not sufficient, for TE insertion.

We might also logically predict the lowest levels of TE silencing activity in the genomes where TEs are most likely to be “on the attack;” however, our results are not fully consistent with this prediction, either. In contrast, based on our analysis of TE-mapping piRNA expression per TE expression, our “TE-attacking” genomes actually exhibit the highest piRNA mapping per TE transcript in the ovaries, but low to intermediate piRNA mapping per TE in the testes. None of the TE-silencing pathways are expressed at consistently lower levels in the “on the attack” genomes, nor are there consistently fewer TE-mapping piRNAs or TE contigs that are mapped by TEs.

In contrast to the three genomes where TEs are most likely to be “on the attack,” all of which are on the larger end of our sampled range of genome size, the three genomes in which TEs are likely to be “on the defense” (based on static or decreasing genome size) span ∼30 Gb of genome size. The largest − *A. davidianus*, with a genome size of 50 pg − shows some patterns consistent with having the lowest TE activity (three apparently inactive TE superfamilies and no superfamilies currently at a historic high, only half as many recognizable (and therefore somewhat recent) LTR/Gypsy genomic contigs as *R. sibiricus* despite having a much higher percentage of LTR/Gypsy sequences in the genome). In addition, *A. davidianus* shows some patterns consistent with high levels of TE silencing: high piRNA pathway expression, numbers of TE-mapping piRNAs, and TRIM28 in the ovary (Figure 5D). However, other patterns (e.g. relatively high levels of TE expression per genomic abundance, relatively low levels of piRNA pathway gene expression in the testes, relatively low levels of KRAB domain expression in both sexes) are inconsistent with well-silenced TEs in a genome maintaining a constant size. Similarly, *T. verrucosus* − with a genome size of 24.5 pg that is likely static or decreasing − shows a mix of patterns associated with degrees of TE activity and silencing: relatively high numbers of unique piRNAs mapping to TEs in both the male and female gonads, but also relatively high TE expression per genomic abundance.

Taken together, these patterns lead us to hypothesize that, once lineages achieve gigantic genome sizes, the long-term outcome of TE/host antagonistic coevolution is largely stochastic TE accumulation. In this scenario, TE abundance is less determined by specific attributes of the proliferation potential of the TEs themselves or of the silencing capacity of the hosts, but instead reflects noisy processes at the genomic and intracellular levels. TE silencing relies on physical interactions between TE and host molecules; for example, piRNA-guided RISC complexes interact with TE transcripts in the cytoplasm and with nascent chromatin-associated transcripts in the nucleus (Aravin, et al. 2008; Reuter, et al. 2011; Iwasaki, et al. 2015; Czech, et al. 2018). Because genome size, nucleus size, and cell size are correlated, species with gigantic genomes also have gigantic nuclear and cell volumes, which means that TE/host silencing molecular interactions play out in large and therefore noisy intracellular arenas (Taylor 2023; Taylor, et al. 2024). Thus, the outcome of a host silencing/TE community interaction, even given the same fundamental players on the TE and host sides, may be noisier in larger cells, weakening any long-term evolutionary correlation between attributes of TEs or hosts and overall TE activity. We cannot completely exclude the possibilities that our inferences of genome expansion or contraction are incorrect (as they are based on sparse genome size data), or that our RNA-and small RNA-based measures of TE silencing activity are failing to accurately represent host silencing activity, or that our measures of TE expression are failing to accurately represent TE activity. However, stochastic dynamics of TE accumulation at the genomic and intracellular levels would be expected to yield overall patterns of genome size evolution that are well-fit by models of quantitative trait evolution that include a strong stochastic component, as seen in salamanders (Mueller, et al. 2023). More practically, our in-depth analyses of these six gigantic genomes, spanning a size range equal to ten human genomes, reveal that any pairwise comparisons between an exemplar “big” and “small” genome focused on a subset of characteristics of TE silencing machinery may lead to spurious conclusions.

Overall, our results lead us to propose that the evolutionary forces that shape the exploration of “genome space” (a multidimensional parameter space including many aspects of a genome, analogous to morphospace) are different when considering small versus large genomes. In the parts of genome space characterized by small genome sizes, we predict that lineages that explore new size space would show signatures of some aspect of their TE silencing correlating with the size shift, and we would predict that aspect of TE silencing to be deterministic to the evolution of genome size; for example, a grasshopper lineage that has undergone genome expansion shows under-expression of the piRNA methylase HENMT (Liu, et al. 2022), and a frog lineage that has undergone genome contraction shows copy number expansion of Piwi genes (Lamichhaney, et al. 2021). In contrast, in the parts of genome space characterized by large genome size, we predict that lineages that explore new size space do so as a consequence of a noisy and stochastic variation-generating process emerging from a weakly antagonistic arms race between TEs and host silencing pathways, and that there may not be alterations to TE silencing that are deterministic to even large evolutionary shifts in genome size.

## Materials and methods

### Specimen information

We collected four male and four female adults of each salamander species: *T. verrucosus*, *P. brevipes, C. orientalis*, *P. hongkongensis* (family Salamandridae) and *A. davidianus* (family Cryptobranchidae) in the breeding season in 2017 (Supplementary Table S1). Collection, euthanasia, and dissection were performed following Animal Care & Use Protocols approved by Chengdu Institute of Biology, Chinese Academy of Sciences.

### Genomic shotgun library creation, sequencing, and assembly

Total DNA was extracted from muscle tissue using the modified low-salt CTAB extraction of high-quality DNA procedure (Arseneau 2017). DNA quality and concentration were assessed using agarose gel electrophoresis and a NanoDrop Spectrophotometer (ThermoFisher Scientific, Waltham, MA), and a PCR-free library was prepared using the NEBNext Ultra DNA Library Prep Kit for Illumina. Sequencing was performed on one lane of a HiSeq 2500 platform (PE250). Library preparation and sequencing were performed by the Beijing Novogene Bioinformatics Technology Co. Ltd. Raw reads were quality-filtered and adaptor-trimmed using Trimmomatic-0.39 (Bolger, et al. 2014) with default parameters.

In total, each genomic shotgun dataset included 12,129,476 to 33,153,950 reads. After filtering and trimming, 11,080,362 to 30,343,958 reads covering a total length of 2,403,712,090 to 6,506,579,244 bp remained. Thus, based on the genome sizes we estimated, the sequencing coverage ranges from 6.4% to 14.7% for unmerged reads. Filtered, trimmed reads were assembled into contigs using dipSPAdes 3.12.0 (Bankevich et al., 2012) with default parameters, yielding 166,335 (*T. verrucosus*) to 649,430 (*P. breviceps*) contigs with N50 values ranging from 475 to 626 bp (Supplementary Table S2).

### Mining and classification of repeat elements

The PiRATE pipeline was used to mine and classify repeats as in the original publication (Berthelier, et al. 2018), including the following steps: 1) Contigs representing repetitive sequences were identified from the assembled contigs using similarity-based, structure-based, and repetitiveness-based approaches. The similarity-based detection programs included RepeatMasker v-4.1.0 (http://repeatmasker.org/RepeatMasker/, using Repbase20.05_REPET.embl.tar.gz as the library) and TE-HMMER (Eddy 2011). The structure-based detection programs included LTRharvest (Ellinghaus, et al. 2008), MGEScan non-LTR (Rho and Tang 2009), HelSearch (Yang, et al. 2009), MITE-Hunter (Han and Wessler 2010), and SINE-finder (Wenke, et al. 2011). The repetitiveness-based detection programs included TEdenovo (Flutre, et al. 2011) and RepeatScout (Price, et al. 2005). 2) Repeat consensus sequences (*i.e.*, sequences representing multiple subfamilies within a TE family) were also identified from the cleaned, filtered, and unassembled reads with dnaPipeTE (Goubert, et al. 2015) and RepeatModeler (http://www.repeatmasker.org/RepeatModeler/). 3) Contigs identified by each individual program in steps 1 and 2, above, were filtered to remove those < 100 bp in length and clustered with CD-HIT-est (Li and Godzik 2006) to reduce redundancy (100% sequence identity cutoff). 4) These contigs were annotated as TEs to the levels of order and superfamily in Wicker’s hierarchical classification system (Wicker, et al. 2007), modified to include several recently discovered TE superfamilies using PASTEC (Hoede, et al. 2014), and checked manually to filter chimeric contigs and those annotated with conflicting evidence (Table 1). 5) All classified repeats (“known TEs” hereafter), along with the unclassified repeats (“unknown repeats” hereafter) and putative multi-copy host genes, were combined to produce a repeat library for each salamander species. **7**) For each superfamily, we collapsed the contigs to 95% and 80% sequence identity using CD-HIT-est to provide an overall view of within-superfamily diversity; 80% is the sequence identity threshold used to define TE families (Wicker, et al. 2007).

### Characterization of the overall repeat element landscape

Overlapping paired-end shotgun reads were merged using PEAR v.0.9.11 (Zhang, et al. 2014) with the following parameter values based on our library insert size and trimming parameters: min-assemble-length 36, max-assemble-length 490, min-overlap size 10. After merging, 12,036,739 to 34,237,973 reads remained (including both merged and singletons), with an N50 of 249-362 bp and total length of 2,267,438,309-6,029,429,512 bp. Following merging, the final sequencing coverage ranged from 5.84% to 13.92%. To calculate the percentage of the genome composed of different TEs, trimmed and merged shotgun reads were masked with RepeatMasker v-4.1.0 using two versions of our repeat library: one that included the unknown repeats and the other that excluded them. This comparison provided a rough approximation of the quantity of unknown repeats that were TE-derived, but divergent, fragmented, or otherwise unidentifiable by our pipeline. In both cases, simple repeats were identified using the Tandem Repeat Finder module implemented in RepeatMasker. The overall results were summarized at the levels of TE class, order, and superfamily.

### Estimation of GC content of TE orders

To calculate the GC content of the different TE orders, bowtie2 was used to map the clean reads to our repeat library (excluding unknown repeats and multi-copy genes), allowing us to classify the aligned reads by TE orders. Overall GC content was then calculated for the reads assigned to each order using the seqkit software (Shen, et al. 2016).

### Summary of the amplification history of TE orders to test for ongoing activity

To summarize the overall amplification history of TE orders and test for ongoing activity, the perl script parseRM.pl (Kapusta, et al. 2017) was used to parse the raw output files from RepeatMasker (.align). The proportion of different TE orders in the genome was calculated by parseRM.pl results (parameter-p-v) and the degree of divergence between each read and consensus sequence was reported (parameter-l 50, 1-a). The repeat library used to mask the reads comprised the contigs classified by the PiRATE pipeline and clustered at 100% sequence identity. Each TE order is represented by multiple consensus contigs that represent ancestral sequences likely corresponding to the family and subfamily TE taxonomic levels (i.e., not the distant common ancestor of the entire order). For each order, histograms were plotted to summarize the percent divergence of all reads from their closest (i.e., least divergent) consensus sequence. These histograms do not allow the delineation between different amplification dynamics scenarios (i.e., a single family with continuous activity versus multiple families with successive bursts of activity). Rather, these global overviews were examined for overall shapes consistent with ongoing activity (i.e., the presence of TE loci < 1% diverged from the ancestral sequence and a unimodal, right-skewed, J-shaped, or monotonically decreasing distribution).

### Phylogenetic analysis of genomic sequences of LTR/Gypsy

Contigs annotated as TEs of the abundant superfamily LTR/Gypsy that contained an identifiable reverse transcriptase domain were selected from the genomic assemblies of each of our five focal species as well as *Ranodon sibiricus*. Sequences longer than 300 amino acids were selected and aligned with MAFFT (Katoh and Standley 2013). Positions with data missing from >20% of OTUs were cut from the initial alignment, and unalignable regions were eliminated. Remaining OTU sequences <200 amino acids in length were then removed from analysis, yielding a final alignment of 1,952 sequences (116 from *A. davidianus*, 545 from *C. orientalis*, 463 from *P. brevipes*, 476 from *P. hongkongensis*, 245 from *R. sibiricus*, and 107 from *T, verrucosus*) with an average length of 300 amino acids (minimum = 200, maximum = 322). We estimated the unrooted phylogenetic tree using maximum likelihood implemented in Iqtree (Minh, et al. 2020) with the best-fitting amino acid substitution model (JTT + R10) selected using BIC and bootstrap support assessed with 1000 replicates; tree visualization was performed with ggtree (Yu, et al. 2017). We assessed whether large clades of LTR/Gypsy contigs were present in larger genomes, consistent with TE proliferation in this superfamily contributing to genome expansion; specifically, we focused on the sister taxa of *P. brevipes* (∼40 Gb) and *P. hongkongensis* (∼50 Gb), which diverged ∼15 mya.

### Comparison of genomic TE community diversity across genome sizes

To test for a relationship between genome size and TE diversity, genomic reference contigs of 88 species including non-tetrapod “fishes” (3 cartilaginous fishes, 13 ray-finned fishes, and 3 lobe-finned fishes), amphibians (10 frogs, 11 salamanders including the 5 from this study, and 4 caecilians), non-avian reptiles (8 snakes, 3 lizards, 2 crocodilians, 2 turtle, and the tuatara), 14 birds, and 14 mammals were downloaded from CNCB (China National GeneBank DataBase) and NCBI databases. Repeats in each genome were annotated de novo using RepeatModeler, and these repeats were then used for each species as the RepeatMasker library to characterize the overall repeat element landscape, as detailed above. Percentages of each genome consisting of each TE superfamily were summarized using parseRM (parameter-p) (Kapusta, et al. 2017). Diversity of the overall TE community in each species was calculated using the Shannon index *H*′ = − ∑ *P_i_* ln(*P_i_*) (Shannon 1948) and the Gini-Simpson index 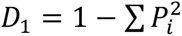, where *P_i_* is the proportion of sequences belonging to TE superfamily *i* (Wang, et al. 2021). Only the top 10 most abundant TE superfamilies were included because of uncertainty in the abundance of rare TE superfamilies. Within each vertebrate clade (salamanders, frogs, birds, mammals, ray-finned fishes) or non-monophyletic group (non-avian reptiles), linear regression analysis was performed to test for a relationship between TE diversity and genome size.

### Estimation of ectopic recombination-mediated deletion of LTR elements

For each salamander species, all genomic contigs > 3,000 bp in length that were annotated to LTR/Gypsy were further annotated using LTRpred to identify terminal and internal sequences (Cho, et al. 2019). Internal and terminal sequences were confirmed by manually checking for internal TE domains using NCBI BLASTx (https://blast.ncbi.nlm.nih.gov/Blast.cgi?PROGRAM=blastx&PAGE_TYPE=BlastSearch&LINK_LOC=blasthome) and for terminal repeat sequences using the NCBI-Blast2suite to align each contig sequence against itself. Internal sequences were conservatively defined to be bounded by the first and last TE domains. Contigs that lacked the complete structure were removed from further analysis. This yielded a total of 12 – 25 LTR/Gypsy contigs per species, with an average terminal sequence length of 408 bp (range 87–971 bp) and an average internal sequence length of 3,541 bp (range 221–6,042 bp). *Tylototriton verrucosus* was excluded from analysis because we only recovered one LTR/Gypsy contig, likely reflecting the lower sequencing coverage. To estimate levels of terminal sequences (LTRs) relative to internal sequences from the remaining four species plus *Ranodon sibiricus* (Wang, et al. 2021), genomic shotgun reads were mapped to the assembled genomic contigs using bowtie2 in local alignment mode with very-sensitive-local preset options and otherwise default parameters, increasing the G-value from the default of 20 to 30, 40, and 50 to increase minimum alignment length for reads (Langmead and Salzberg 2012). This analysis was performed twice: once treating all reads as unpaired and once using merged paired-end reads plus unmerged reads. Average read depths across the terminal and internal portion in each of the selected LTR/Gypsy contigs were estimated by scaling the number of hits by the lengths of the terminal and internal regions. From these estimates, the total terminal-to-internal sequence ratio (T:I) was calculated for each contig. In the absence of ectopic recombination mediated by terminal repeats, this ratio would be 1:1; increasing levels of ectopic recombination would produce ratios > 1:1. We tested for the predicted relationship between deletion rate and genome size across our focal salamander species (i.e larger genomes showing lower deletion rates) using ANOVA.

### Transcriptome library creation, sequencing, assembly, and TE annotation

Total RNA was extracted separately from testes (*n* = 4) and ovary (*n* = 4) tissues using TRIzol (Invitrogen). For each sample, RNA quality and concentration were assessed using agarose gel electrophoresis, a NanoPhotometer spectrophotometer (Implen, CA), a Qubit 2.0 Fluorometer (ThermoFisher Scientific), and an Agilent BioAnalyzer 2100 system (Agilent Technologies, CA), requiring an RNA integrity number (RIN) of 7.5 or higher. Sequencing libraries were generated using the NEBNext Ultra RNA Library Prep Kit for Illumina following the manufacturer’s protocol. After cluster generation of the index-coded samples, the library was sequenced on one lane of an Illumina Hiseq 4000 platform (PE 150). Transcriptome sequences were filtered using Trimmomatic-0.39 with default parameters (Bolger et al., 2014). 28,294,628 to 51,120,646 reads were retained for each testis or ovary sample. Remaining reads of all testes and ovary samples were combined and assembled using Trinity 2.12.0 (Haas et al., 2013), yielding 452,395 to 690,114 contigs (i.e., putative assembled transcripts) (Supplementary Table S11). For each species, assembled contigs were clustered using CD-hit-est (95% identity). Completeness of these final *de novo* transcriptome assemblies were assessed using the BUSCO pipeline with both the vertebrata and tetrapoda BUSCO datasets (Simao et al., 2015). For all five assemblies, the BUSCO completeness score was >94% using the vertebrata dataset and >87% using the tetrapoda dataset.

Expression levels of contigs in each sample were measured with Salmon (Patro et al., 2017), and contigs with no raw counts were removed. To annotate the remaining contigs containing autonomous TEs, BLASTp and BLASTx were used against the Repbase Database (downloaded on January 5, 2022) with an E-value cutoff of 1E-5 and 1E-10, respectively. The aligned length coverage was set to exceed 80% of the queried transcriptome contigs. To annotate contigs containing non-autonomous TEs, RepeatMasker was used with our genomic repeat library of non-autonomous TEs (LARD-, TRIM-, MITE-, and SINE-annotated contigs) and the requirement that the transcriptome/genomic contig overlap was > 80 bp long, > 80% identical in sequence, and covered > 80% of the length of the genomic contig. Contigs annotated as conflicting autonomous and non-autonomous TEs were filtered out.

To identify contigs that contained endogenous genes, the Trinotate annotation suite (Bryant et al., 2017) was used; BLASTp and BLASTx were used against the Uniprot database with an E-value cutoff of 1E-5 and 1E-10, respectively, and 1E-5 was used as the cutoff for the HMMER search against the Pfam database (Wheeler and Eddy, 2013). To identify contigs that contained both a TE and an endogenous gene (i.e., putative cases where a TE and a gene were co-transcribed on a single transcript), all contigs that were annotated both by Repbase and Trinotate were examined, and the ones annotated by Trinotate to contain a TE-encoded protein (*i.e.*, the contigs where Repbase and Trinotate annotations were in agreement) were not further considered. The remaining contigs annotated by Trinotate to contain a non-TE gene (*i.e.*, an endogenous salamander gene) and also annotated either by Repbase to include a TE-encoded protein or by RepeatMasker to include a non-autonomous TE were identified for further examination and expression-based analysis.

### Germline TE expression quantification in males and females

Expression levels of contigs in each sample were measured with Salmon (Patro et al., 2017). For each species, the male and female expression levels of each TE-annotated contig were calculated by averaging the TPM values among replicates of each sex. Next, the across-replicate average TPMs of all contigs annotated to each superfamily were summed for males and for females, yielding TE superfamily expression levels for each sex. For TE superfamilies detected in both the genomic and transcriptomic datasets, we tested for a relationship between genomic abundance (in base pairs, calculated as percentage of the total genome size occupied by each superfamily) and expression levels in each sex using linear regression on log-transformed data. We tested whether the males and females had significantly different slopes or intercepts for each species, and we also tested for across-species differences in slope for each sex. To identify differentially expressed contigs between males and females, DESeq2 (Love et al., 2014) was used with an adjusted P-value cutoff of 0.01 and a minimal log_2_fold-change cutoff of 2, and the contigs representing TEs were identified. For each sex in each species, we also summarized the average percentage of total expression (measured as TPM) for TEs, endogenous protein-coding genes, and unannotated contigs (including both unknown repeats as well as contigs that received no annotation). We assessed whether any of these relationships (i.e. TE expression relative to genomic abundance, differential TE expression between sexes) changed with increasing genome size.

### Identification of putative piRNAs from small RNA-seq data

Small RNA libraries were prepared for each sample using the NEBNext® Multiplex Small RNA Library Prep Set for Illumina® (NEB, USA) following the manufacturer’s recommendations (Wang, et al. 2021). Library quality was assessed on the Agilent Bioanalyzer 2100 using DNA High Sensitivity Chips. Clustering of index-coded samples was performed on a cBot Cluster Generation System using the TruSeq SR Cluster Kit v3-cBot-HS (Illumina) according to the manufacturer’s instructions. After cluster generation, libraries were sequenced on an Illumina Hiseq 2500 platform (SE50). Libraries for *P. hongkongensis* ovaries failed for technical reasons and the samples were excluded from further analysis.

Low-quality sequences were filtered using the fastq_quality_filter (-q 20,-p 90) in the FASTX-Toolkit v0.0.13 (http://hannonlab.cshl.edu/fastx_toolkit/). Adapter sequences were removed with a minimum overlap of 10 bp from the 3′-end, untrimmed reads were discarded, and those with a minimum length of 18 bp, a maximum length of 40 bp (after cutting adapters), and no Ns were selected using cutadapt v2.8 (Martin 2011). Reads mapping to the mitochondrial genomes (NCBI accession numbers: *T. verrucosus* MF461428, *P. brevipes* NC_053711*, C. orientalis* KY399474, *P. hongkongensis* AY458597, and *A. davidianus* KT119359) and riboRNAs (NCBI accession numbers: DQ283664, AJ279506, MH806872) were identified and filtered out using Bowtie v1.1.0 (Langmead et al., 2009). Finally, we filtered out any sequences appearing only once in the dataset as putative sequencing artifacts. Overall, more reads were filtered out based on length and rRNA identity in females than males (Supplementary Table S12). We conservatively defined putative piRNAs as those ranging from 25-30 nt, and we selected these using seqkit (Shen et al., 2016). For each individual, we calculated the number of unique piRNA sequences per 10,000,000 clean reads 18-40 nt in length.

### Putative piRNAs targeting TE superfamilies

To estimate levels of piRNAs targeting each TE superfamily, putative piRNAs were mapped to the transcriptome assembly using bowtie with default parameter settings, and piRNAs that map to autonomous TEs (i.e., those that include transposition-related ORFs) in the sense and antisense orientations were identified. Reads per million (RPM) values were calculated for each TE contig and then averaged across individuals of each sex. For each sex, overall putative piRNA levels targeting each TE superfamily were calculated by summing across all contigs annotated to the same TE superfamily. We tested for a relationship between TE superfamily expression levels and levels of mapping/targeting piRNAs in each sex using linear regression on log-transformed data. We tested whether the males and females had significantly different slopes or intercepts for each species, and we also tested for across-species differences in slope for each sex. In addition, we quantified the average percentage of TE contigs that were mapped by piRNAs in both sexes. We assessed whether the relationship between TEs and their targeting piRNAs changed with increasing genome size.

### Ping-pong piRNA biogenesis signature analysis

piRNA biogenesis associated with piRNA-targeted post-transcriptional TE silencing produces a distinctive “ping-pong signature” in the piRNA pool, which consists of a 10 bp overlap between the 5′ ends of antisense and sense piRNAs. The ping-pong signature for each individual was analyzed using the following approach: First, TE transcripts that were not mapped by both sense-oriented and antisense-oriented piRNAs were filtered out using Bowtie, allowing 0 mismatches for sense mapping under the assumption that piRNAs derived directly from an RNA target should have the identical sequence (Teefy et al., 2020) and 3 mismatches for antisense mapping because cleavage of RNA targets can occur with imperfect base-pairing (Zhang et al., 2015). Second, the fractions of overlapping pairs of sense/antisense piRNAs corresponding to specific lengths, as well as the Z-score measuring the significance of each ping-pong signature, were generated using the 1_piRNA_and_Degradome_Counts.RMD and 3_Ping_Pong_Phasing. Rmd scripts (Teefy et al., 2020). We calculated the average values (fraction overlapping, Z-score) for females and males for each species, and we assessed whether the signature of ping-pong amplification changed with increasing genome size for either testes or ovaries.

### Query for piRNA cluster transcripts

piRNAs are processed from much longer precursor transcripts of genomic loci known as piRNA clusters, which include numerous active and inactive transposable element insertions (Ku and Lin 2014). In an attempt to identify putative primary piRNA cluster transcripts, putative piRNAs (i.e. 25-30 nt in length) from each individual were mapped to the respective species’ transcriptome assembly using srnaMapper (Zytnicki and Gaspin 2022) and the mapping results were input into proTRAC (Rosenkranz and Zischler 2012) to identify piRNA cluster transcripts. piRNA cluster identification is typically performed using these tools to map smallRNA-seq data to a genome assembly, but both approaches can also be used with a transcriptome assembly instead without violating any assumptions. Because our libraries are poly-A selected, our analysis would only identify cluster transcripts that have undergone polyadenylation. Contigs identified as putative piRNA cluster transcripts were annotated for TE insertions as well as protein-coding genes, and the percentage of total putative piRNAs mapping to putative cluster transcripts was calculated. Expression levels of piRNA cluster contigs in each sample were measured with Salmon (Patro et al., 2017). For each species, the male and female expression levels of each cluster-annotated contig were calculated by averaging the TPM values among replicates of each sex.

### Germline TE silencing pathway expression across genome sizes

We identified transcripts of 22 genes receiving a direct annotation of piRNA processing in vertebrates in the Gene Ontology knowledgebase that were present in the majority of our target species: ASZ1, BTBD18 (BTBDI), DDX4, EXD1, FKBP6, GPAT2, HENMT1 (HENMT), MAEL, MOV10l1 (M10L1), PIWIL1, PIWIL2, PIWIL4, PLD6, PNLDC1 (PNDC1), TDRD1, TDRD5, TDRD6, TDRD7, TDRD9, TDRD12 (TDR12), TDRD15 (TDR15), and TDRKH. In addition, we identified transcripts of 16 genes encoding proteins that create a transcriptionally repressive chromatin environment in response to recruitment by PIWI proteins or KRAB-ZFP proteins, 16 of which received a direct annotation of NuRD complex in the Gene Ontology knowledgebase and 2 of which were taken from the literature: CBX5, CHD3, CHD4, CSNK2A1, DNMT1, GATAD2A (P66A), HDAC1, MBD3, MTA1, MTA2, MTA3, RBBP4, RBBP7, SALL1, SETDB1 (SETB1), and ZBTB7A (ZBT7A) (Wang, et al. 2021)(Ecco et al., 2017). Additionally, we identified TRIM28, which bridges this repressive complex to TE-bound KRAB-ZFP proteins in lobe-finned fishes (Ecco et al., 2017). For comparison, we identified transcripts of 16 protein-coding genes receiving a direct annotation of miRNA processing in vertebrates in the Gene Ontology knowledgebase, which we did not predict to differ in expression based on genome size: DSRAD, AGO1, AGO2, AGO3, AGO4, DICER1, NUP155 (NU155), PUM1, PUM2, RBM3, SNIP1, SPOUT1 (CI114), TARBP2 (TRBP2), TRIM71 (LIN41), ZC3H7A, and ZC3H7B. Finally, we identified all transcripts annotated with a KRAB domain in each species. Expression levels for each transcript in each individual were measured with Salmon (Patro et al., 2017). RBM3 was removed from the *A. davidianus* analysis because its expression was several orders of magnitude higher than in any of the other species.

As a proxy for overall piRNA silencing activity, for each individual, we calculated the ratio of total piRNA pathway expression (summed TPM of 22 genes) to total miRNA pathway expression (summed TPM of 16 genes). As a proxy for transcriptional repression driven by both the piRNA pathway and KRAB-ZFP binding activity, we calculated the ratio of total transcriptional repression machinery expression (summed TPM of 16 genes) to total miRNA pathway expression. Additionally, we calculated the ratio of TRIM28 expression to total miRNA pathway expression for each individual. Finally, we calculated the summed TPM of all transcripts annotated with a KRAB domain in each individual. We tested for differences between males and females within each species using t-tests, and we tested for differences across species in males and in females using ANOVA. We assessed whether overall piRNA silencing activity, transcriptional repression, TRIM28 expression, or KRAB domain expression in either sex changed with increasing genome size.

## 5 Data availability statement

Genomic shotgun and transcriptome sequences have been deposited in the Genome Sequence Archive at the National Genomics Data Center, Beijing Institute of Genomics, Chinese Academy of Sciences/China National Center for Bioinformation (GSA: CRA******, CRA******, CRA******), and are publicly accessible at http://bigd.big.ac.cn/gsa.

## 6 Author contributions

**Jie Wang** & **Rachel Lockridge Mueller**: Conceptualization, Formal analysis, Investigation, Resources, Writing-review & editing, Supervision, Project administration. All authors read and approved the final manuscript. **Guangpu Zhang, Cheng Sun, Guiying Chen, Michael W. Itgen & Ava Haley**: Methodology, Software. **Liming Chang**, **Yingyong Wang & Xin Yang**: Specimen collection. **Jiaxing Tang**: Methodology, Formal analysis, Writing-original draft, Visualization. All authors read and approved the final manuscript.

## 7 Funding

This work was supported by the National Natural Science Foundation of China (Grant Nos. 32170435, 31570391), the Emergency Open Competition Project of National Forestry and Grassland Administration (202303), and the National Science Foundation of the United States of America (Grant No. 1911585 to RLM).

## Acknowledgements

We gratefully acknowledge Wancai Xia and Lusha Liu for target tissue dissection, and Jianping Jiang, Jiongyu Liu and Weiye Deng for expertise in facilitating data analysis.

## 8 Conflict of Interest

The authors declare that the research was conducted in the absence of any commercial or financial relationships that could be construed as a potential conflict of interest.

## Notes

### Competing Interest Statement

The authors have declared no competing interest.

