## Supplementary Files for "Size evolution of gigantic genomes suggests stochastic outcomes of transposable element/host silencing interactions"

### Supplementary Materials

#### S1 Samples and data used in this study.

| Species | Tissue | Code | Sex | Mass: g | STL: cm | Collection date | Collection sites | Source | Genomic sequencing | | RNA quality | | | | mRNA sequencing | | small RNA sequencing | |
|---|---|---|---|---|---|---|---|---|---|---|---|---|---|---|---|---|---|---|
| | | | | | | | | | Raw reads | Accession code[1] | OD260/280 | OD260/230 | 28S/18S | RIN | Raw reads | Accession code[2] | Raw reads | Accession code[3] |
| *Ranodon sibiricus* | Testis#1 | WQ07 | ♂ | 16.83 | 17.252 | 2017.8.22. | Wenquan County, Xinjiang Uygur Autonomous Region | wild | | | 1.77 | 2.75 | 1.7 | 9.6 | 37,888,360 | CRR610084 | 19,717,749 | CRR610091 |
| | Testis#2 | WQ08 | ♂ | 19.79 | 18.192 | 2017.8.22. | | wild | | | 1.73 | 0.95 | 1.1 | 8.9 | 35,386,396 | CRR610085 | 21,815,001 | CRR610092 |
| | Testis#3 | WQ10 | ♂ | 12.02 | 16.441 | 2017.8.22. | | wild | | | 2.00 | 3.22 | 1.7 | 9.4 | 34,854,594 | CRR610086 | 16,177,680 | CRR610093 |
| | Testis#4 | WQ05 | ♂ | 20.48 | 15.591 | 2017.8.22. | | captive | | | 1.70 | 2.13 | 1.7 | 9.3 | 31,732,246 | CRR610083 | 19,005,099 | CRR610090 |
| | Ovary#1 | WQ03 | ♀ | 32.85 | 21.001 | 2017.8.22. | | captive | 11,960,858 | CRR609958 | 1.96 | 0.68 | 1.1 | 9.3 | 38,466,202 | CRR610080 | 17,965,762 | CRR610088 |
| | Ovary#2 | WQ02 | ♀ | 28.61 | 20.804 | 2017.8.22. | | captive | | | 1.64 | 0.49 | 1.5 | 9.6 | 40,660,732 | CRR610079 | 15,836,634 | CRR610087 |
| | Ovary#3 | WQ04 | ♀ | 35.37 | 20.437 | 2017.8.22. | | captive | | | 1.64 | 0.22 | 0.8 | 8.7 | 39,418,784 | CRR610081 | 14,063,357 | CRR610089 |
| | Ovary#4 | WQ06 | ♀ | 23.57 | 18.149 | 2017.8.22. | | captive | | | 1.66 | 0.15 | 0.1 | 7.7[4] | 39,229,748 | CRR610082 | | |
| *Tylototriton verrucosus* | Testis#1 | HS02 | ♂ | 13.0 | 14.1 | 2017.7.8. | Husa County, Yunnan province | Wild | | | 1.99 | 2.31 | 1.6 | 9.3 | 32,691,262 | CRR649922 | 16,585,370 | CRR649930 |
| | Testis#2 | HS04 | ♂ | 13.2 | 15.0 | 2017.7.8. | | Wild | | | 1.94 | 2.42 | 1.7 | 9.2 | 32,973,554 | CRR649923 | 17,619,593 | CRR649931 |
| | Testis#3 | HS05 | ♂ | 13.0 | 15.2 | 2017.7.8. | | Wild | | | 1.99 | 2.01 | 1.7 | 9.6 | 42,699,602 | CRR649924 | 17,707,096 | CRR649932 |
| | Testis#4 | HS07 | ♂ | 20.2 | 15.4 | 2017.7.8. | | Wild | | | 2.02 | 2.32 | 1.6 | 9.3 | 35,485,366 | CRR649925 | 16,278,837 | CRR649933 |
| | Ovary#1 | HS08 | ♀ | 25.8 | 17.5 | 2017.7.8. | | Wild | | | 1.58 | 0.25 | 0.6 | 6.5 | 39,101,474 | CRR649918 | 31,842,294 | CRR649926 |
| | Ovary#2 | HS09 | ♀ | 34.3 | 20.2 | 2017.7.8. | | Wild | 12,129,476 | CRR649917 | 1.84 | 0.97 | 1.1 | 8.5 | 39,718,882 | CRR649919 | 18,122,932 | CRR649927 |
| | Ovary#3 | HS11 | ♀ | 25.0 | 18.6 | 2017.7.8. | | Wild | | | 1.79 | 0.73 | 0.9 | 7.8 | 38,441,274 | CRR649920 | 17,009,880 | CRR649928 |
| | Ovary#4 | HS12 | ♀ | 20.1 | 16.7 | 2017.7.8. | | Wild | | | 1.61 | 0.29 | 0.6 | 7.8 | 40,016,632 | CRR649921 | 35,999,752 | CRR649929 |

| | | | | | | | | | | | | | | | | | | |
|---|---|---|---|---|---|---|---|---|---|---|---|---|---|---|---|---|---|---|
| | Testis#1 | NF01 | ♂ | 9.8 | 13.1 | 2017.7.9. | Nanfeng | wild | | | 2.00 | 2.22 | 1.1 | 9.4 | 38,735,632 | CRR649888 | 17,289,840 | CRR649896 |
| | Testis#2 | NF14 | ♂ | 31.5 | 20.0 | 2017.7.9. | County, | wild | | | 1.91 | 2.32 | 1.4 | 9.8 | 37,515,790 | CRR649889 | 18,259,136 | CRR649897 |
| | Testis#3 | NF16 | ♂ | 29.3 | 18.6 | 2017.7.9. | Jiangxi | wild | | | 2.0 | 2.18 | 1.4 | 9.7 | 35,913,846 | CRR649890 | 19,728,618 | CRR649898 |
| *Pachytriton* | Testis#4 | NF18 | ♂ | 15.9 | 15.4 | 2017.7.9. | Province | wild | | | 1.95 | 2.27 | 1.3 | 9.7 | 41,001,140 | CRR649891 | 18,206,032 | CRR649899 |
| *brevipes* | Ovary#1 | NF08 | ♀ | 17.0 | 15.3 | 2017.7.9. | | wild | | | 1.58 | 0.35 | 0.9 | 8.5 | 32,094,588 | CRR649884 | 32,733,395 | CRR649892 |
| | Ovary#2 | NF09 | ♀ | 16.5 | 15.8 | 2017.7.9. | | wild | | | 1.95 | 1.45 | 1.5 | 8.3 | 34,827,584 | CRR649885 | 18,218,210 | CRR649893 |
| | Ovary#3 | NF20 | ♀ | 22.3 | 18.3 | 2017.7.9. | | wild | | | 1.99 | 1.41 | 1.0 | 8.7 | 33,516,658 | CRR649886 | 17,778,912 | CRR649894 |
| | Ovary#4 | NF21 | ♀ | 21.7 | 17.4 | 2017.7.9. | | wild | 15,505,860 | CRR650566 | 1.95 | 2.22 | 1.5 | 8.3 | 40,024,958 | CRR649887 | 18,167,875 | CRR649895 |
| | Testis#1 | CO16 | ♂ | 1.2 | 6.9 | 2017.10.17. | Kecheng | wild | | | 1.50 | 0.33 | 1.2 | 8.3 | 41,446,178 | CRR649563 | 17,538,653 | CRR649571 |
| | Testis#2 | CO17 | ♂ | 1.2 | 6.0 | 2017.10.17. | District, | wild | | | 1.54 | 0.95 | 1.0 | 8.1 | 33,512,742 | CRR649564 | 19,588,629 | CRR649572 |
| | Testis#3 | QZ01 | ♂ | 1.8 | 6.3 | 2017.7.4. | Quzhou | wild | | | 1.67 | 1.88 | 0.7 | 7.6 | 32,573,566 | CRR649565 | 15,203,225 | CRR649573 |
| *Cynops* | Testis#4 | QZ04 | ♂ | 2.0 | 6.7 | 2017.7.4. | City, | wild | | | 1.96 | 1.93 | 0.7 | 8.1 | 41,743,244 | CRR649566 | 18,690,834 | CRR649574 |
| *orientalis* | Ovary#1 | CO12 | ♀ | 2.1 | 7.6 | 2017.10.17. | Zhejiang | wild | | | 1.86 | 1.68 | 0.8 | 8.5 | 34,189,104 | CRR649559 | 16,659,083 | CRR649567 |
| | Ovary#2 | CO13 | ♀ | 2.0 | 8.1 | 2017.10.17. | Province | wild | | | 1.90 | 1.95 | 0.6 | 6.7 | 34,644,206 | CRR649560 | 18,525,552 | CRR649568 |
| | Ovary#3 | CO14 | ♀ | 1.6 | 7.5 | 2017.10.17. | | wild | | | 1.90 | 1.83 | 0.5 | 7.4 | 54,097,378 | CRR649561 | 15,023,175 | CRR649569 |
| | Ovary#4 | QZ09 | ♀ | 3.2 | 7.6 | 2017.7.4. | | wild | 33,153,950 | CRR649558 | 1.81 | 0.62 | 0.5 | 8.3 | 36,944,040 | CRR649562 | 16,661,516 | CRR649570 |
| | Testis#1 | LY06 | ♂ | | 74.0 | 2017.8.10. | Lueyang | captive | | | 1.86 | 1.84 | 1.2 | 8.6 | 29,719,324 | CRR638172 | 16,008,097 | CRR638180 |
| | Testis#2 | LY07 | ♂ | | 86.0 | 2017.8.10. | County, | captive | | | 1.87 | 1.77 | 2.3 | 8.2 | 34,960,348 | CRR638173 | 17,458,603 | CRR638181 |
| | Testis#3 | LY08 | ♂ | | 87.5 | 2017.8.10. | Shaanxi | captive | | | 2.04 | 1.09 | 0.9 | 7.2 | 53,487,888 | CRR638174 | 19,122,025 | CRR638182 |
| *Andrias* | Testis#4 | LY09 | ♂ | | 89.5 | 2017.8.10. | Province | captive | | | 1.88 | 1.56 | 1.2 | 8.7 | 33,144,322 | CRR638175 | 20,191,827 | CRR638183 |
| *davidianus* | Ovary#1 | LY01 | ♀ | | 71.5 | 2017.8.10. | | captive | 24,305,964 | CRR638167 | 1.00 | 0.23 | 1.5 | 8.2 | 31,801,468 | CRR638168 | 17,631,197 | CRR638176 |
| | Ovary#2 | LY03 | ♀ | | 79.0 | 2017.8.10. | | captive | | | 1.97 | 2.25 | 0.9 | 7.0 | 32,312,166 | CRR638169 | 18,539,178 | CRR638177 |
| | Ovary#3 | LY04 | ♀ | | 71.0 | 2017.8.10. | | captive | | | 1.99 | 2.40 | 0.9 | 7.5 | 34,674,288 | CRR638170 | 16,260,101 | CRR638178 |
| | Ovary#4 | LY05 | ♀ | | 75.0 | 2017.8.10. | | captive | | | 1.96 | 2.44 | 0.8 | 7.3 | 37,821,156 | CRR638171 | 18,605,559 | CRR638179 |

| | | | | | | | | | | | | | | | | | | |
|---|---|---|---|---|---|---|---|---|---|---|---|---|---|---|---|---|---|---|
| *Paramesotriton honkongensis* | Testis#1 | HK03 | ♂ | 7.5 | 12.0 | 2017.12.8. | Shenzhen City, Guangdong Province | wild | | | 1.82 | 0.68 | 1.2 | 9.0 | 41,780,356 | CRR649905 | 16,326,142 | CRR649913 |
| | Testis#2 | HK04 | ♂ | 7.4 | 11.5 | 2017.12.8. | | wild | | | 1.82 | 0.98 | 0.9 | 8.3 | 39,777,108 | CRR649906 | 18,469,923 | CRR649914 |
| | Testis#3 | HK05 | ♂ | 8.6 | 12.0 | 2017.12.8. | | wild | | | 1.68 | 0.68 | 1.0 | 8.2 | 39,191,664 | CRR649907 | 16,805,777 | CRR649915 |
| | Testis#4 | HK06 | ♂ | 9.4 | 11.5 | 2017.12.8. | | wild | | | 1.80 | 1.06 | 0.9 | 8.5 | 41,074,192 | CRR649908 | 18,347,342 | CRR649916 |
| | Ovary#1 | HK08 | ♀ | 10.6 | 14.0 | 2017.12.8. | | wild | | | 1.51 | 0.21 | 0.4 | 5.9 | 37,576,072 | CRR649901 | 9,308,482 | CRR649909 |
| | Ovary#2 | HK09 | ♀ | 8.2 | 11.5 | 2017.12.8. | | wild | | | 1.49 | 0.15 | 0.5 | 7.4 | 38,208,494 | CRR649902 | 15,027,311 | CRR649910 |
| | Ovary#3 | HK10 | ♀ | 10.6 | 12.5 | 2017.12.8. | | wild | | | 1.38 | 0.17 | 0.4 | 7.0 | 42,629,786 | CRR649903 | 21,879,203 | CRR649911 |
| | Ovary#4 | HK11 | ♀ | 10.3 | 12.0 | 2017.12.8. | | wild | 16,943,516 | CRR649900 | 1.58 | 0.22 | 0.9 | 8.3 | 35,881,698 | CRR649904 | 18,743,046 | CRR649912 |

[1]https://ngdc.cncb.ac.cn/gsa/s/5d3dHx5W

[2]https://ngdc.cncb.ac.cn/gsa/s/p15v402F

[3]https://ngdc.cncb.ac.cn/gsa/s/X3i83677

#### S2 Steps for the genomic assembly and the estimation of sequencing depth.

| Steps | Types | *Ranodon sibiricus* WQ03 | *Tylototriton verrucosus* HS09 | *Pachytriton brevipes* NF21 | *Cynops orientalis* QZ09 | *Andrias davidianus* LY01 | *Paramesotriton honkongensi* HK11 |
|---|---|---|---|---|---|---|---|
| | Raw reads | 11,960,858 | 12,129,476 | 15,505,860 | 33,153,950 | 24,305,964 | 16,943,516 |
| Trimmomatic | Clean reads | 11,168,678 | 11,080,362 | 14,218,520 | 30,343,958 | 22,113,929 | 14,752,452 |
| | N50 of clean reads | 250 | 246 | 250 | 245 | 243 | 250 |
| | Sum length of clean reads | 2,314,096,923 | 2,403,712,090 | 3,195,482,437 | 6,506,579,244 | 4,634,122,878 | 3,222,403,582 |
| Sequencing Depth | Genome size[#] | 21,300,000,000 | 23,961,000,000 | 38,924,400,000 | 43,325,400,000 | 48,900,000,000 | 49,878,000,000 |
| | Depth | 0.11 X | 0.10 X | 0.08 X | 0.15 X | 0.09 X | 0.06 X |
| Merged by Pear | No. of merged pair-reads | 3,783,512 | 9,518,030 | 9,870,941 | 25,315,531 | 18,112,904 | 10,935,902 |
| | Unassembled forward/reverse reads | 1,500,762 | 3,661,719 | 2,165,798 | 8,922,442 | 6,256,605 | 2,750,468 |
| | N50 of reads | 388 | 250 | 362 | 250 | 249 | 250 |
| | Sum length of reads | 1,997,175,501 | 2,267,438,309 | 2,787,991,555 | 6,029,429,512 | 4,256,895,753 | 2,911,584,397 |
| Assembled by dipSpades | No. of contigs | 478,991 | 166,335 | 649,430 | 662,111 | 276,974 | 634,076 |
| | Sum length of contigs | 249,425,929 | 95,192,733 | 360,507,191 | 429,174,065 | 161,443,731 | 368,170,024 |
| | Min length of contigs | 128 | 128 | 128 | 130 | 128 | 128 |
| | N50 of contigs | 447 | 536 | 475 | 626 | 540 | 516 |
| | Max length of contigs | 25,462 | 16,789 | 20,021 | 18,448 | 28,984 | 26,186 |

[#] Data from Gregory 2024. Animal genome size database. http://www.genomesize.com.

**S3.1 Classification of repeat contigs (modified from Wicker 2007) and summary of repeats detected in the genome of *Ranodon sibiricus* (J. Wang et al., 2023).**

| Order | Superfamily | Percent of Genome[a] | Genomic Contigs (100% Identical) | Genomic Contigs (95% Identical) | Genomic Contigs (80% Identical) | Average Genomic Contig Length (100% identical) (bp) | Longest Genomic Contig (bp) | Transcriptome contigs (80% Identical) | Average Expression Level in females (TPM) | Average Expression Level in males (TPM) |
|---|---|---|---|---|---|---|---|---|---|---|
| **Class I - Retrotransposons - Autonomous** | | | | | | | | | | |
| LTR | *Gypsy* | 3.85-6.50 | 5,985 | 5,464 | 3,835 | 548 | 7,587 | 2,057 | 827 | 3,024 |
| | *ERV* | 0.41-0.43 | 413 | 394 | 264 | 593 | 11,567 | 321 | 392 | 694 |
| | *Copia* | 0.10-0.25 | 23 | 21 | 19 | 592 | 1,353 | 27 | 31 | 26 |
| | *Bel-Pao* | 0.05 | 12 | 7 | 2 | 575 | 766 | - | - | - |
| | *Retrovirus* | - | 3 | 2 | 2 | 1,375 | 1,719 | 10 | 1 | 10 |
| | *THE1* | - | 4 | 4 | 3 | 491 | 674 | - | - | - |
| | Unknown LTR | - | 4 | 4 | 4 | 469 | 955 | - | - | - |
| DIRS | *DIRS* | 4.44-5.95 | 8,087 | 7,821 | 3,844 | 379 | 4,266 | 4,844 | 5,427 | 11.962 |
| PLE | *Penelope* | 0.09-0.12 | 482 | 462 | 376 | 407 | 3,208 | 460 | 123 | 352 |
| LINE | *Jockey* | 9.69-12.12 | 25,276 | 23,361 | 13,287 | 426 | 3,625 | 11,622 | 6,206 | 15,535 |
| | *L1* | 5.04-6.62 | 10,189 | 8,997 | 5,941 | 672 | 6,546 | 8,241 | 2,954 | 7,610 |
| | *RTE* | 0.12-0.24 | 227 | 201 | 137 | 676 | 4,540 | 399 | 130 | 360 |
| | *I* | 0.09-0.17 | 32 | 25 | 10 | 905 | 3,858 | 48 | 42 | 125 |
| | *R2* | - | | | | | | 1 | - | 1 |
| | Unknown LINE | 0.39-1.89 | 90 | 77 | 72 | 1,393 | 5,881 | 1 | - | - |
| **Class I - Retrotransposons - Non-autonomous** | | | | | | | | | | |
| SINE | *5S* | 0.23-0.18 | 57 | 41 | 15 | 773 | 1,783 | 4 | 5 | 1 |
| | *7SL* | - | 43 | 41 | 15 | 243 | 450 | 2 | 1 | - |
| | *tRNA* | - | 22 | 22 | 22 | 273 | 403 | - | - | - |
| | Unknown SINE | 0.45-3.42 | 365 | 348 | 338 | 377 | 695 | 219 | 2,901 | 2,471 |

| | | | | | | | | | | |
|---|---|---|---|---|---|---|---|---|---|---|
| Retrotransposon | TRIM | 3.80-9.76 | 748 | 631 | 465 | 588 | 2,619 | 492 | 1,651 | 5,847 |
| Derivatives | LARD | 0.15-0.73 | 45 | 37 | 35 | 2,834 | 8,148 | 153 | 515 | 2,643 |
| **Class II DNA transposons Subclass 1** | | | | | | | | | | |
| | *PIF-Harbinger* | 2.98-4.22 | 1,164 | 1,014 | 434 | 411 | 5,169 | 416 | 575 | 1,359 |
| | *hAT* | 1.15-1.12 | 177 | 135 | 55 | 869 | 5,706 | 35 | 83 | 26 |
| | *Tc1-Mariner* | 0.18-0.63 | 73 | 40 | 24 | 1025 | 3,781 | 20 | 58 | 87 |
| | *PiggyBac* | 0.05-0.08 | 9 | 8 | 6 | 1,087 | 1,956 | 6 | 21 | 15 |
| | *MuDR* | 0 | 3 | 3 | 3 | 266 | 417 | 13 | 4 | 10 |
| TIR | *CACTA* | 0 | 2 | 2 | 2 | 645 | 846 | 3 | 2 | 1 |
| | *ISL2EU* | - | | | | | | 1 | 18 | 16 |
| | *Ginger* | - | | | | | | 4 | 10 | 6 |
| | *Academ* | - | | | | | | 11 | 5 | 8 |
| | *P* | - | | | | | | 1 | 1 | 1 |
| | Unknown TIR | 0.33-1.46 | 22 | 20 | 19 | 754 | 2,174 | - | - | - |
| Transposon Derivatives | MITE | 0.56-4.07 | 357 | 346 | 321 | 258 | 1,942 | 137 | 465 | 1,188 |
| **Class II - DNA Transposons - Subclass 2** | | | | | | | | | | |
| Maverick | *Maverick* | 0.05 | 258 | 228 | 65 | 583 | 6,090 | 123 | 44 | 762 |
| Helitron | *Helitron* | 0.18-0.48 | 49 | 38 | 19 | 939 | 6,551 | 13 | 2 | 10 |
| **Total** | | 34.38-60.54 | 54,221 | 49,794 | 29,634 | 489 | 11,567 | 29,684 | 22,494 | 42,200 |

[a]The first and second numbers were estimated including and excluding unknown repeats, respectively, from the repeat library.

#### S3.2 Classification of repeat contigs (modified from Wicker 2007) and summary of repeats detected in the genome of *Tylototriton verrucosus*

| Order | Superfamily | Percent of Genome (%)[a] | Genome Contigs (100% identical) | genome contigs (95% identical) | genome contigs (80% identical) | Average genome contig length (100% iedntical) | Longest genomic contig | transcriptome contigs (80% identical) | Average Expression Level in females (TPM) | Average Expression Level in males (TPM) |
|---|---|---|---|---|---|---|---|---|---|---|
| Class I Retrotransposon-Autonomous | | | | | | | | | | |
| LTR | Gypsy | 7.83~14.06 | 2,736 | 2,463 | 1,629 | 523 | 7,664 | 4614 | 5978 | 24646 |
| | ERV | 1.55~2.03 | 746 | 568 | 307 | 493 | 3,198 | 824 | 1255 | 11480 |
| | Copia | 0 | 1 | 1 | 1 | 214 | 214 | 87 | 416 | 0 |
| | Retrovirus | 0.71~0.90 | 38 | 20 | 6 | 1,082 | 5,401 | 48 | 406 | 1631 |
| | Unknown LTR | 0.01~0.02 | 4 | 4 | 4 | 443 | 584 | 1 | 3 | 0 |
| DIRS | DIRS | 2.07~2.80 | 2,352 | 2,223 | 1,330 | 465 | 5,232 | 5369 | 5524 | 48717 |
| PLE | Penelope | 0.13~0.14 | 344 | 343 | 309 | 314 | 2,161 | 1392 | 1082 | 3604 |
| LINE | Jockey | 2.13~2.45 | 1,350 | 1,173 | 694 | 460 | 3,173 | 2350 | 3038 | 16417 |
| | L1 | 2.79~3.03 | 4,998 | 4,584 | 2,897 | 692 | 4,057 | 7016 | 5210 | 22665 |
| | RTE | 0.15~0.16 | 197 | 177 | 158 | 691 | 2,640 | 1082 | 736 | 3523 |
| | I | 0 | 1 | 1 | 1 | 122 | 122 | 15 | 32 | 54 |
| | R2 | 0 | 2 | 2 | 2 | 1,072 | 1,419 | 2 | 0 | 9 |
| | Unknown LINE | 0.09~0.15 | 7 | 6 | 5 | 978 | 1,392 | - | - | - |
| **Class I Retrotransposon Non-autonomous** | | | | | | | | | | |
| SINE | 5S | 0.04 | 1 | 1 | 1 | 851 | 851 | 2 | 4 | 70 |
| | Alu | 0.01 | 25 | 24 | 11 | 218 | 360 | 3 | 107 | 80 |
| | Unknown SINE | 0.36~1.00 | 80 | 67 | 60 | 380 | 4,309 | 84 | 259 | 2096 |
| Retrotransposon derivative | TRIM | 1.99~3.22 | 108 | 72 | 54 | 635 | 1,666 | 132 | 4749 | 10838 |
| | LARD | 0.08~0.18 | 11 | 9 | 9 | 2,223.90 | 3,099 | 17 | 146 | 112 |

| | | | | | | | | | | |
|---|---|---|---|---|---|---|---|---|---|---|
| **unknown ClassI** | unknown ClassI | 0 | 2 | 2 | 2 | 585 | 952 | - | - | - |
| **Class II DNA transposons Subclass 1** | | | | | | | | | | |
| TIR | hAT | 0.06~0.13 | 44 | 41 | 36 | 479 | 2,144 | 105 | 1487 | 1295 |
| | Tc1-Mariner | 0 | 22 | 22 | 22 | 413 | 1,153 | 15 | 232 | 68 |
| | Kolobok | - | - | - | - | - | - | 7 | 3098 | 981 |
| | PiggyBac | 0.14~0.18 | 21 | 16 | 6 | 895 | 2,323 | 19 | 57 | 562 |
| | PIF-Harbinger | 5.04~12.52 | 467 | 328 | 90 | 552 | 4,467 | 240 | 2095 | 6541 |
| | CACTA | - | - | - | - | - | - | 5 | 8 | 25 |
| | MuDR | 0 | 4 | 4 | 3 | 326 | 534 | 12 | 65 | 54 |
| | Ginger | 0 | 2 | 2 | 2 | 421 | 705 | 14 | 65 | 203 |
| | P | 0 | 1 | 1 | 1 | 299 | 299 | - | - | - |
| | Academ | 0 | 8 | 8 | 6 | 590 | 2,287 | 20 | 52 | 304 |
| | Sola | 0 | 2 | 1 | 1 | 224 | 231 | - | - | - |
| | Novosib | 0.12~0.45 | 29 | 28 | 23 | 356 | 833 | | | |
| | Zisupton | - | - | - | - | - | - | 3 | 0 | 4 |
| | Unknown TIR | 0.1~0.11 | 181 | 179 | 140 | 402 | 1,207 | - | - | - |
| Crypton | Crypton | 0.1~0.11 | 21 | 21 | 19 | 1,770 | 5,955 | 1 | 3 | 1 |
| Transposon derivative | MITE | 0.4~2.96 | 122 | 114 | 104 | 234 | 2,679 | 53 | 369 | 2805 |
| **Class II DNA transposon Subclass 2** | | | | | | | | | | |
| Maverick | Maverick | 0.14 | 73 | 55 | 22 | 771 | 5,661 | 77 | 158 | 2244 |
| Helitron | Helitron | 0.03 | 15 | 14 | 7 | 383 | 935 | 32 | 4 | 107 |
| unknown ClassII | *unknown Class*II | 0 | 2 | 2 | 2 | 379 | 516 | - | - | - |
| Total | | 26.07 ~ 46.82 | 14,006 | 12,576 | 7,964 | 18,712 | 77,324 | 23,641 | 36,638 | 161,136 |

[a]The first and second numbers were estimated including and excluding unknown repeats, respectively, from the repeat library.

#### S3.3 Classification of repeat contigs (modified from Wicker 2007) and summary of repeats detected in the genome of *Pachytriton brevipes*

| Order | Superfamily | Percent of Genome[a] | Genomic Contigs(100 %Identical) | Genomic Contigs (95%Identical) | Genomic Contigs (80%Identical) | Average Genomic Contig Length (100% identical) | Longest Genomic Contig | Transcriptome contigs (80% Identical) | Average Expression Level in females(TPM) | Average Expression Level in males (TPM) |
|---|---|---|---|---|---|---|---|---|---|---|
| **Class I - Retrotransposons - Autonomous** | | | | | | | | | | |
| LTR | *Gypsy* | 5.10 ~ 7.83 | 14,331 | 12,967 | 7,721 | 474.9 | 6,483 | 2534 | 3314 | 14381 |
| | *ERV* | 2.25 ~ 2.74 | 4,241 | 2,778 | 873 | 431.1 | 10,078 | 578 | 1782 | 11090 |
| | *Copia* | 0.17 ~ 0.38 | 22 | 22 | 20 | 541.5 | 5,265 | 1 | 0 | 76 |
| | *Retrovirus* | 0.76 ~ 0.95 | 40 | 31 | 18 | 1,086 | 3,261 | 64 | 166 | 1855 |
| | *Unknown LTR* | 0.23 ~ 0.49 | 13 | 13 | 12 | 515.8 | 2,296 | 2 | 3 | 2 |
| DIRS | *DIRS* | 3.41 ~ 4.89 | 10,137 | 9,447 | 5,011 | 428.2 | 7,363 | 3099 | 3409 | 40461 |
| PLE | *Penelope* | 0.78 ~ 1.44 | 3,127 | 3,075 | 2,384 | 337 | 6,401 | 688 | 376 | 1647 |
| LINE | *Jockey* | 4.57 ~ 6.47 | 11,590 | 10,610 | 5,571 | 474.6 | 3,601 | 2519 | 2535 | 15714 |
| | *L1* | 4.10 ~ 6.24 | 20,570 | 19,481 | 13,743 | 515 | 3,696 | 6489 | 3477 | 16085 |
| | *RTE* | 0.55 ~ 0.77 | 1,444 | 1,303 | 899 | 653.1 | 3,521 | 670 | 433 | 1309 |
| | *I* | 0 | 4 | 4 | 3 | 191.8 | 261 | 6 | 0 | 8 |
| | *R2* | 0 | 2 | 2 | 2 | 912.5 | 1,482 | 2 | 62 | 273 |
| | *Unknown LINE* | 0.07 ~ 1.19 | 5 | 5 | 5 | 1,076 | 3,385 | - | - | - |
| **Class I - Retrotransposons - Non-autonomous** | | | | | | | | | | |
| SINE | *5S* | 0 | 7 | 7 | 5 | 124.3 | 151 | 3 | 18 | 144 |
| | *Alu* | 0.05 ~ 0.06 | 79 | 72 | 35 | 246.7 | 678 | 2 | 14 | 20 |
| | *Unknown SINE* | 0.42 ~ 1.22 | 175 | 171 | 134 | 198.2 | 615 | 27 | 100 | 637 |
| Retrotransposon Derivatives | TRIM | 9.38 ~ 11.4 | 216 | 198 | 152 | 710.4 | 4,327 | 187 | 2585 | 12485 |
| | LARD | 0.89 ~ 1.69 | 36 | 36 | 35 | 2,364 | 7,468 | 131 | 329 | 3100 |
| unknown ClassI | unknown ClassI | 0.02 ~ 0.03 | 19 | 18 | 16 | 383.6 | 1,237 | 7 | 1 | 5 |
| **Class II - DNA Transposons - Subclass 1** | | | | | | | | | | |

| | | | | | | | | | | |
|---|---|---|---|---|---|---|---|---|---|---|
| TIR | *PIF-Harbinger* | 1.43 ~ 1.89 | 571 | 463 | 172 | 341 | 4,168 | 92 | 1542 | 1172 |
| | *hAT* | 0.34 ~ 0.40 | 196 | 145 | 76 | 511.9 | 3,366 | 81 | 2968 | 1541 |
| | *Tc1-Mariner* | 0 | 12 | 12 | 12 | 282.8 | 406 | 2 | 127 | 18 |
| | *PiggyBac* | 0.05 | 32 | 31 | 10 | 477.3 | 1,617 | 11 | 239 | 187 |
| | *MuDR* | 0.06 | 30 | 28 | 18 | 480.9 | 3,449 | 22 | 17 | 163 |
| | *Sola* | 0 | 5 | 5 | 5 | 338.2 | 783 | - | - | - |
| | *CACTA* | - | - | - | - | - | - | 1 | 0 | 2 |
| | *Kolobok* | - | - | - | - | - | - | 5 | 4170 | 1619 |
| | *Ginger* | 0 | 1 | 1 | 1 | 1,554 | 1,554 | 7 | 99 | 101 |
| | *Academ* | 0 | 5 | 5 | 5 | 422.6 | 550 | 12 | 9 | 68 |
| | *Zator* | - | - | - | - | - | - | 1 | 0 | 1 |
| | *Novosib* | 0.12 ~ 0.19 | 77 | 76 | 61 | 308.1 | 1,729 | - | - | - |
| | *Unknown TIR* | 0.25 ~ 0.51 | 617 | 614 | 544 | 399.5 | 3,538 | - | - | - |
| Crypton | *Crypton* | 0.43 ~ 0.45 | 142 | 128 | 105 | 1625 | 2,931 | 3 | 1 | 12 |
| Transposon Derivatives | MITE | 0.44 ~ 1.52 | 399 | 385 | 387 | 188.2 | 704 | 60 | 340 | 971 |
| **Class II - DNA Transposons - Subclass 2** | | | | | | | | | | |
| Maverick | *Maverick* | 0.06 | 149 | 111 | 31 | 502.7 | 3,875 | 27 | 67 | 633 |
| *Helitron* | *Helitron* | *0.51 ~ 0.66* | *162* | 126 | 55 | *528.6* | *4,613* | 69 | *81* | *393* |
| **unknown ClassII** | **unknown ClassII** | *0* | 1 | 1 | 1 | 267 | 267 | 1 | 32 | 4 |
| **Total** | | 36.44 ~ 53.58 | 68,457 | 62,371 | 38,122 | 19,892 | 105,119 | 17,403 | 28,296 | 126,177 |

[a]The first and second numbers were estimated including and excluding unknown repeats, respectively, from the repeat library.

#### S3.4 Classification of repeat contigs (modified from Wicker 2007) and summary of repeats detected in the genome of *Cynops orientalis*

| Order | Superfamily | Percent of Genome [a] | Genomic Contigs (100%Identical) | Genomic Contigs (95%Identical) | Genomic Contigs (80%Identical) | Average Genomic Contig Length (100% identical) | Longest Genomic Contig | Transcriptome contigs (80% Identical) | Expression Level in females(TPM) | Expression Level in males(TPM) |
|---|---|---|---|---|---|---|---|---|---|---|
| Class I Retrotransposon-Autonomous | | | | | | | | | | |
| LTR | Gypsy | 6.67 ~ 10.56 | 11,859 | 11,377 | 6,530 | 513.8 | 6,918 | 5568 | 4845 | 16979 |
| | ERV | 1.33 ~ 2.04 | 1,120 | 1,030 | 502 | 504.3 | 9,189 | 868 | 668 | 6594 |
| | Copia | 0.05 ~ 0.08 | 39 | 37 | 32 | 320.1 | 2,791 | 2 | 0 | 3 |
| | Retrovirus | 0.67 ~ 0.76 | 56 | 37 | 16 | 1362.9 | 3,039 | 70 | 126 | 885 |
| | Unknown LTR | 0.40 ~ 0.49 | 3 | 2 | 2 | 619.3 | 1,087 | 1 | 0 | 10 |
| DIRS | DIRS | 3.85 ~ 4.89 | 9.941 | 9,660 | 4,682 | 446.6 | 7,546 | 5033 | 4342 | 38368 |
| PLE | Penelope | 0.44 ~ 0.54 | 2,102 | 2,099 | 1,635 | 280.3 | 5,686 | 2249 | 1429 | 4574 |
| LINE | Jockey | 4.30 ~ 5.48 | 11,475 | 10,802 | 5,183 | 483.4 | 3,359 | 4426 | 3417 | 20134 |
| | L1 | 5.33 ~ 6.05 | 28,897 | 27,360 | 17,278 | 571 | 3,696 | 13797 | 6087 | 30695 |
| | RTE | 0.47 ~ 0.54 | 1,337 | 1,131 | 760 | 800.7 | 3,261 | 1390 | 867 | 2770 |
| | I | 0 | 1 | 1 | 1 | 232 | 232 | 9 | 33 | 17 |
| | R2 | 0 | 2 | 2 | 1 | 1,439.50 | 2,547 | 5 | 187 | 73 |
| | Unknown LINE | 0.04 ~ 0.09 | 9 | 9 | 9 | 832.8 | 1,830 | 5 | 5 | 4 |
| Class I Retrotransposon Non-autonomous | | | | | | | | | | |
| SINE | 5S | 0 | 10 | 10 | 7 | 120.1 | 153 | - | - | - |
| | 7SL | 0 | 1 | 1 | 1 | 302 | 302 | 1 | 12 | 0 |
| | Alu | 0 | 8 | 8 | 7 | 226.9 | 378 | 3 | 43 | 39 |
| | Unknown SINE | 0.33 ~ 1.09 | 171 | 164 | 123 | 223.1 | 1,695 | 58 | 236 | 723 |
| Retrotransposon derivative | TRIM | 4.7 ~ 8.37 | 344 | 217 | 183 | 710.3 | 4,455 | 297 | 3524 | 13636 |
| | LARD | 0.49 ~ 1.45 | 58 | 54 | 52 | 2424.8 | 7,089 | 186 | 539 | 1475 |
| unknown ClassI | unknown ClassI | 0.01 | 33 | 31 | 29 | 308.8 | 1,084 | 19 | 2 | 42 |

| | | | | | | | | | | |
|---|---|---|---|---|---|---|---|---|---|---|
| Class II DNA transposon Subclass 1 | | | | | | | | | | |
| TIR | hAT | 0.57 ~ 0.66 | 119 | 93 | 69 | 746.9 | 6,593 | 113 | 1596 | 721 |
| | Tc1-Mariner | 0 | 11 | 11 | 11 | 351.5 | 765 | 7 | 127 | 16 |
| | Kolobok | - | - | - | - | - | - | 8 | 3978 | 801 |
| | PiggyBac | 0.05 ~ 0.06 | 31 | 26 | 7 | 712.5 | 2,861 | 16 | 273 | 125 |
| | PIF-Harbinger | 2.02 ~ 2.96 | 645 | 507 | 177 | 447.5 | 5,785 | 145 | 331 | 837 |
| | CACTA | - | - | - | - | - | - | 3 | 2 | 6 |
| | MuDR | 0.02 ~ 0.04 | 18 | 18 | 15 | 298.2 | 630 | 23 | 6 | 62 |
| | Ginger | 0 | 1 | 1 | 1 | 1,554 | 1,554 | 12 | 24 | 34 |
| | Academ | 0 | 6 | 6 | 6 | 393.8 | 785 | 12 | 17 | 35 |
| | ISL2EU | - | - | - | - | - | - | 1 | 49 | 2 |
| | Sola | 0 | 1 | 1 | 1 | 250 | 250 | - | - | - |
| | Zisupton | - | - | - | - | - | - | 1 | 0 | 0 |
| | Novosib | 0.02 ~ 0.03 | 33 | 32 | 29 | 291.8 | 959 | - | - | - |
| | Zator | - | - | - | - | - | - | 2 | 0 | 3 |
| | Unknown TIR | 0.19 ~ 0.29 | 1,091 | 1,090 | 900 | 358 | 2,054 | - | - | - |
| Crypton | Crypton | 0.49 ~ 0.54 | 143 | 135 | 106 | 1,799.70 | 4,658 | 6 | 0 | 11 |
| Transposon derivative | MITE | 0.75 ~ 3.01 | 435 | 428 | 407 | 196 | 982 | 92 | 363 | 1268 |
| Class II DNA transposon Subclass 2 | | | | | | | | | | |
| Maverick | Maverick | 0.07 ~ 0.08 | 118 | 106 | 42 | 572.7 | 3,187 | 37 | 69 | 684 |
| Helitron | Helitron | 0.44 ~ 0.69 | 105 | 85 | 46 | 739.7 | 4,280 | 69 | 78 | 249 |
| unknown ClassII | unknown ClassII | 0 | 4 | 4 | 4 | 199.3 | 372 | - | - | - |
| Total | | 33.7 ~ 50.8 | 60,296 | 66,575 | 38,854 | 21,634 | 102,052 | 34,534 | 33,275 | 141,875 |

[a]The first and second numbers were estimated including and excluding unknown repeats, respectively, from the repeat library.

#### S3.5 Classification of repeat contigs (modified from Wicker 2007) and summary of repeats detected in the genome of *Andrias davidianus*

| Order | Superfamily | Percent of Genome[a] | Genomic Contigs(100 %Identical) | Genomic Contigs (95%Identical) | Genomic Contigs (80%Identical) | Average Genomic Contig Length (100%identical) | Longest Genomic Contig | Transcriptome contigs (80% Identical) | Expression Level in females(TPM) | Expression Level in males (TPM) |
|---|---|---|---|---|---|---|---|---|---|---|
| **Class I - Retrotransposons – Autonomous** | | | | | | | | | | |
| | Gypsy | 14.43 ~23.37 | 9,757 | 9,092 | 5,360 | 466.6 | 7,584 | 2467 | 4,209 | 16,675 |
| | ERV | 0.45 ~ 0.58 | 353 | 347 | 262 | 501.6 | 6,764 | 324 | 409 | 1,974 |
| LTR | Retrovirus | 0.01 | 2 | 2 | 2 | 1334.5 | 2,490 | 20 | 23 | 57 |
| | Copia | 0 | 4 | 4 | 4 | 524 | 628 | 75 | 284 | 3 |
| | Unknown LTR | 0 | 3 | 3 | 3 | 167 | 244 | 1 | 0 | 3 |
| DIRS | DIRS | 3.10 ~ 4.94 | 4,836 | 4,746 | 2,103 | 384.5 | 4,348 | 3679 | 57,717 | 69,660 |
| PLE | Penelope | 0.51 ~ 1.27 | 1,491 | 1,478 | 1,213 | 342 | 5,891 | 1318 | 2,458 | 4,377 |
| | Jockey | 4.57 ~ 5.83 | 8,867 | 8,579 | 5,193 | 371.4 | 3,674 | 4978 | 15,839 | 26,889 |
| | L1 | 3.27 ~ 3.95 | 5,162 | 4,989 | 3,359 | 443.1 | 6,610 | 4664 | 4,049 | 17,989 |
| | RTE | 0.09 ~ 0.32 | 70 | 69 | 57 | 482 | 5,365 | 144 | 218 | 514 |
| LINE | I | 0 | 7 | 7 | 6 | 474.3 | 1,253 | 17 | 32 | 68 |
| | R2 | 0 | 2 | 2 | 2 | 527 | 816 | 28 | 414 | 197 |
| | Unknown LINE | 0.03 ~ 0.04 | 2 | 2 | 2 | 852 | 864 | - | - | - |
| Class I - Retrotransposons - Non-autonomous | | | | | | | | | | |
| | 5S | 0 | 1 | 1 | 1 | 128 | 128 | 1 | 4 | 81 |
| | 7SL | 0 | 1 | 1 | 1 | 363 | 363 | - | - | - |
| SINE | Alu | 0.01 | 39 | 38 | 19 | 229.8 | 727 | 11 | 49 | 174 |
| | SVA | | | | | | | 1 | 3 | 0 |
| | Unknown SINE | 0.31 ~ 0.94 | 99 | 98 | 93 | 239.2 | 649 | 73 | 300 | 893 |
| | TRIM | 4.46 ~ 6.19 | 138 | 112 | 65 | 672.9 | 5,511 | 1505.18 | 1,505 | 5,651 |

| | | | | | | | | | | |
|---|---|---|---|---|---|---|---|---|---|---|
| Retrotransposon Derivatives | LARD | 0.34 ~ 1.57 | 20 | 20 | 20 | 3084.5 | 7,373 | 109 | 369 | 1,608 |
| Unknown class I | unknown ClassI | 0 ~ 0.02 | 7 | 5 | 4 | 191.7 | 325 | 12 | 23 | 23 |
| Class II - DNA Transposons - Subclass 1 | | | | | | | | | | |
| TIR | PIF-Harbinger | 0.89 ~ 1.42 | 413 | 380 | 132 | 318.2 | 3,892 | 129 | 322 | 709 |
| | hAT | 0.33 ~ 0.44 | 81 | 71 | 45 | 337.6 | 1,134 | 80 | 630 | 625 |
| | Tc1-Mariner | 0.10 ~ 0.11 | 50 | 35 | 22 | 296.1 | 688 | 40 | 424 | 254 |
| | Novosib | 0.01 ~ 0.04 | 18 | 18 | 17 | 268.8 | 372 | - | - | - |
| | PiggyBac | 0.01 | 7 | 7 | 5 | 467.6 | 1,333 | 8 | 23 | 43 |
| | Academ | 0.01 | 6 | 6 | 5 | 520.3 | 910 | 7 | 36 | 133 |
| | Kolobok | - | - | - | - | - | - | 5 | 2056 | 1186 |
| | CACTA | - | - | - | - | - | - | 2 | 3 | 0 |
| | MuDR | 0 | 1 | 1 | 1 | 471 | 471 | 16 | 78 | 39 |
| | Ginger | 0 | 1 | 1 | 1 | 538 | 538 | 2 | 6 | 2 |
| | P | 0 | 1 | 1 | 1 | 305 | 305 | - | - | - |
| | Unknown TIR | 0.37 ~ 0.63 | 112 | 110 | 78 | 424.3 | 2,576 | - | - | - |
| Crypton | Crypton | 0.21 ~ 0.25 | 29 | 29 | 23 | 1,513.40 | 2,216 | - | - | - |
| Transposon Derivatives | MITE | 0.63 ~ 4.26 | 228 | 227 | 218 | 195 | 1,158 | 91 | 2121 | 3243 |
| Class II - DNA Transposons - Subclass 2 | | | | | | | | | | |
| Maverick | Maverick | 0.16 ~ 0.26 | 159 | 148 | 89 | 553.7 | 3,829 | 67 | 470 | 586 |
| Helitron | Helitron | 0.06 ~ 0.08 | 16 | 15 | 11 | 522.9 | 2,362 | 11 | 8 | 4,894 |
| unknown ClassII | unknown ClassII | 0.01 ~ 0.01 | 17 | 8 | 6 | 190.8 | 403 | 1 | - | - |
| Total | | 34.37~56.56 | 32,000 | 30,652 | 18,423 | 18,702 | 83,794 | 18,381 | 94,082 | 158,550 |

[a]The first and second numbers were estimated including and excluding unknown repeats, respectively, from the repeat library.

#### S3.6 Classification of repeat contigs (modified from Wicker 2007) and summary of repeats detected in the genome of *Paramesotriton honkongensis*

| Order | Superfamily | Percent of Genome[a] | Genomic Contigs (100%Identical) | Genomic Contigs (95%Identical) | Genomic Contigs (80%Identical) | Average Genomic Contig Length (100% identical) | Longest Genomic Contig | Transcriptome contigs (80% Identical) | Expression Level in females (TPM) | Expression Level in males(TPM) |
|---|---|---|---|---|---|---|---|---|---|---|
| **Class I - Retrotransposons - Autonomous** | | | | | | | | | | |
| LTR | *Gypsy* | 7.27~10.28 | 12,700 | 12,251 | 8,347 | 473 | 11,581 | 6,431 | 8,420 | 25,662 |
| | *ERV* | 3.17~4.69 | 2,227 | 1,958 | 994 | 512 | 12,607 | 1,008 | 4,091 | 15,182 |
| | *Copia* | 0.05~0.08 | 6 | 6 | 6 | 372 | 1,409 | 31 | 72 | 119 |
| | *Retrovirus* | 1.19~1.59 | 70 | 50 | 18 | 1,186.00 | 3,158 | 92 | 997 | 3,135 |
| | *Unknown LTR* | 0.14~0.71 | 11 | 11 | 11 | 1,378 | 6,092 | 4 | 3 | 9 |
| DIRS | *DIRS* | 2.97~3.79 | 7,501 | 7,335 | 4,481 | 434 | 7,171 | 6,359 | 22,104 | 80,955 |
| PLE | *Penelope* | 1.12~1.32 | 4,343 | 4,328 | 3,397 | 313.9 | 5,670 | 3,570 | 2,654 | 9,016 |
| LINE | *Jockey* | 3.71~4.38 | 9,514 | 9,000 | 5,195 | 488 | 3,542 | 6,352 | 9,650 | 29,090 |
| | *L1* | 4.06~4.56 | 16,670 | 16,090 | 12,100 | 511 | 7,355 | 19,197 | 14,948 | 52,747 |
| | *RTE* | 0.66~0.79 | 1,450 | 1,284 | 943 | 715 | 4,583 | 1,867 | 1,572 | 4,879 |
| | *I* | 0 | 4 | 4 | 4 | 235 | 291 | 21 | 9 | 47 |
| | *R2* | 0.02 | 7 | 6 | 1 | 827 | 4,100 | 3 | 14 | 64 |
| | *Unknown LINE* | 0.06~0.46 | 7 | 5 | 5 | 715 | 4,583 | 3 | 2 | 2 |
| **Class I - Retrotransposons - Non-autonomous** | | | | | | | | | | |
| SINE | *5S* | 0.01~0.02 | 11 | 9 | 3 | 413 | 1,735 | - | - | - |
| | *7SL* | 0 | 1 | 1 | 1 | 196 | 196 | - | - | - |
| | *Alu* | 0.01 | 28 | 28 | 13 | 247 | 444 | 4 | 1 | 23 |
| | *Unknown SINE* | 0.21~1.04 | 169 | 166 | 132 | 214 | 1,071 | 39 | 508 | 1778 |
| Retrotransposon Derivatives | TRIM | 5.35~7.77 | 296 | 206 | 155 | 805 | 4,354 | 291 | 8,332 | 17,011 |
| | LARD | 0.66~1.85 | 67 | 64 | 61 | 2,277.00 | 6,205 | 190 | 875 | 2,219 |
| **unknown ClassI** | unknown ClassI | 0~0.01 | 19 | 19 | 19 | 291 | 949 | | | |

| | | | | | | | | | | |
|---|---|---|---|---|---|---|---|---|---|---|
| **Class II - DNA Transposons - Subclass 1** | | | | | | | | | | |
| TIR | *PIF-Harbinger* | 2.09~3.19 | 524 | 430 | 167 | 523 | 12,770 | 133 | 767 | 1545 |
| | *hAT* | 0.63~0.98 | 187 | 150 | 100 | 851 | 6,459 | 93 | 1,690 | 1,050 |
| | *Tc1-Mariner* | 0 | 17 | 17 | 17 | 376 | 1,153 | 5 | 141 | 28 |
| | *PiggyBac* | 0.08~0.09 | 46 | 42 | 16 | 561 | 2,768 | 14 | 162 | 236 |
| | *MuDR* | 0.05~0.06 | 22 | 22 | 15 | 375 | 1,418 | 28 | 36 | 132 |
| | *CACTA* | - | - | - | - | - | - | 16 | 29 | 52 |
| | *Kolobok* | - | - | - | - | - | - | 9 | 3,941 | 1,624 |
| | *Ginger* | 0 | 8 | 8 | 6 | 419 | 1,554 | 10 | 161 | 43 |
| | *Academ* | 0 | 17 | 17 | 17 | 401 | 1,332 | 19 | 35 | 47 |
| | *P* | 0 | 571 | 473 | 184 | 526 | 12,770 | - | - | - |
| | *Zator* | - | - | - | - | - | - | 1 | 2 | 10 |
| | *Novosib* | 0.09~0.13 | 72 | 71 | 65 | 287 | 957 | - | - | - |
| | *Unknown TIR* | 0.18~0.28 | 490 | 488 | 425 | 342 | 1,922 | - | - | - |
| Crypton | *Crypton* | 0.50~0.53 | 137 | 128 | 95 | 1,729 | 3,078 | 10 | 17 | 9 |
| Transposon Derivatives | MITE | 0.54~1.88 | 381 | 378 | 371 | 178 | 727 | 100 | 356 | 1,053 |
| **Class II - DNA Transposons - Subclass 2** | | | | | | | | | | |
| Maverick | *Maverick* | 0.09~0.10 | 127 | 119 | 51 | 625 | 3,188 | 39 | 136 | 1,098 |
| *Helitron* | *Helitron* | *1.63~2.33* | *237* | 174 | *64* | 899 | *10,888* | 104 | *221* | *890* |
| **unknown ClassII** | *unknown Class*II | *0.47~1.44* | *4* | *3* | *3* | *442* | *528* | - | - | - |
| **Total** | | 37.01~54.38 | 57,371 | 54,869 | 37,299 | 20,913 | 136,140 | 46,043 | 81,946 | 249,755 |

[a]The first and second numbers were estimated including and excluding unknown repeats, respectively, from the repeat library.

**S4 GC content of TE superfamilies across salamander species.**

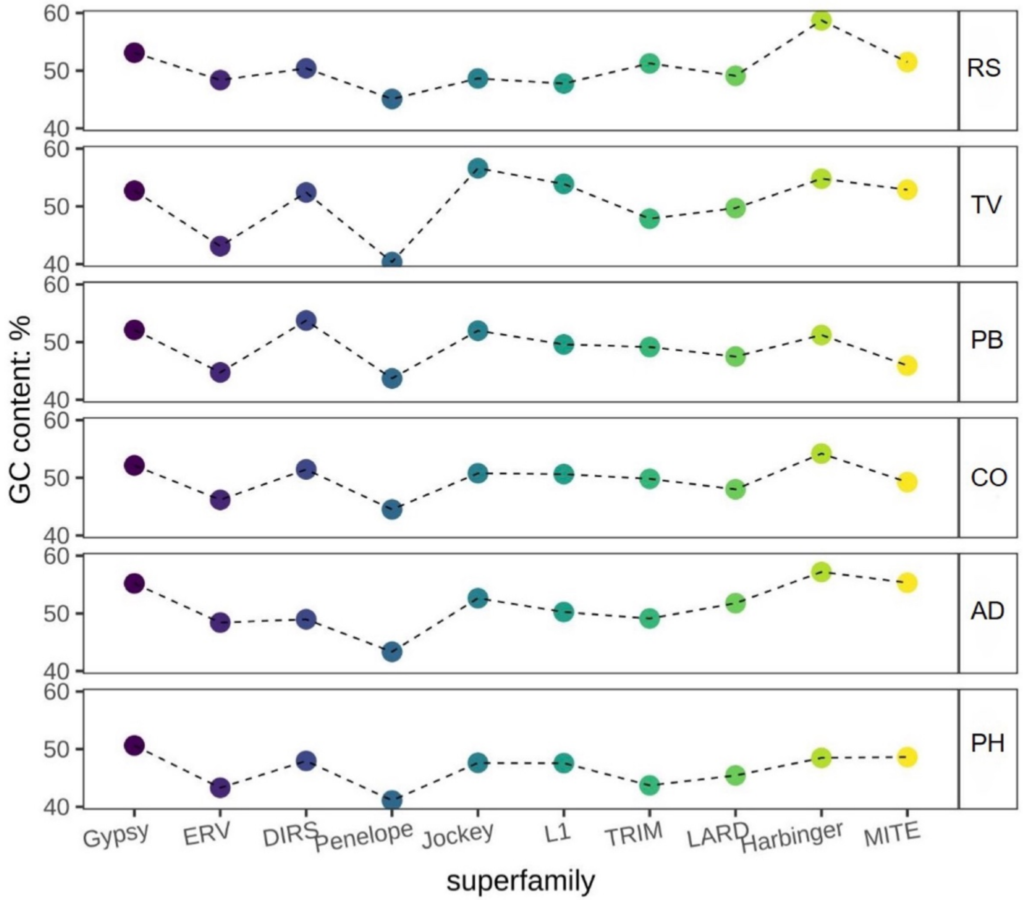

Species are arranged top to bottom by increasing genome size: RS, *Ranodon sibiricus*; PB, *Pachytriton brevipes*; CO, *Cynops orientalis*; AD, *Andrias davidianus*; PH, *Paramesotriton hongkongensis*.

**S5 Shannon diversity index (SI) and Gini-Simpson diversity index (GS) summarizing the TE communities of 88 vertebrate species with different genome size (GS). The six focal salamander species are in bold and red.**

| Chinese name | English_name | species | Family | GS | SI | GSI | Type |
|---|---|---|---|---|---|---|---|
| 菊黄东方鲀 | yellowbelly pufferfish | *Takifugu flavidus* | Tetraodontidae | 0.38 | 2.01 | 0.83 | Bony Fishes |
| 红鳍东方鲀 | fugu | *Takifugu rubripes* | Tetraodontidae | 0.38 | 1.95 | 0.81 | Bony Fishes |
| 半滑舌鳎 | Chinese tongue sole | *Cynoglossus semilaevis* | Cynoglossidae | 0.47 | 1.56 | 0.69 | Bony Fishes |
| 莱茵河杜父鱼 | Rheingroppe | *Cottus rhenanus* | Cottidae | 0.56 | 1.92 | 0.80 | Bony Fishes |
| 太平洋蓝鳍金枪鱼 | North Pacific bluefin tuna | *Thunnus orientalis* | Scombridae | 0.83 | 1.83 | 0.78 | Bony Fishes |
| 白鲟 | white sturgeon | *Acipenser transmontanus* | Acipenseridae | 0.86 | 2.06 | 0.85 | Bony Fishes |
| 雀鳝 | Alligator gar | *Atractosteus spatula* | Lepisosteidae | 1.10 | 1.99 | 0.82 | Bony Fishes |
| 鲤 | common carp | *Cyprinus carpio* | Cyprinidae | 1.68 | 2.15 | 0.87 | Bony Fishes |
| 斑马鱼 | zebrafish | *Danio rerio* | Cyprinidae | 1.70 | 2.11 | 0.85 | Bony Fishes |
| 金线鲃 | Barbus grahami | *Sinocyclocheilus grahami* | Cyprinidae | 1.75 | 2.09 | 0.86 | Bony Fishes |
| 俄罗斯鲟鱼 | Russian sturgeon | *Acipenser gueldenstaedtii* | Acipenseridae | 2.40 | 2.11 | 0.86 | Bony Fishes |
| 大马哈鱼 | Dog salmon | *Oncorhynchus keta* | Salmonidae | 2.60 | 1.70 | 0.75 | Bony Fishes |
| 大西洋鲑鱼 | Atlantic salmon | *Salmo salar* | Salmoninae | 3.00 | 1.79 | 0.77 | Bony Fishes |
| 腔棘鱼 | African coelacanth | *Latimeria chalumnae* | Latimeriidae | 2.74 | 1.90 | 0.81 | Lobe-finned fish |
| 大白鲨 | white shark | *Carcharodon carcharias* | Alopiidae | 4.63 | 1.53 | 0.66 | Cartilaginous fish |
| 点纹斑竹鲨 | brownbanded bamboo shark | *Chiloscyllium punctatum* | Hemiscylliidae | 4.70 | 1.31 | 0.58 | Cartilaginous fish |
| 云纹猫鲨 | cloudy catshark | *Scyliorhinus torazame* | Scyliorhinidae | 6.70 | 1.74 | 0.77 | Cartilaginous fish |
| 非洲肺鱼 | African lungfish | *Protopterus annectens* | Protopteridae | 40.00 | 1.63 | 0.72 | Lobe-finned fish |
| 澳大利亚肺鱼 | Australian lungfish | *Neoceratodus forsteri* | Ceratodontidae | 43.00 | 1.45 | 0.69 | Lobe-finned fish |
| 加蓬鱼螈 | Gaboon caecilian | *Geotrypetes seraphini* | Dermophiidae | 3.80 | 1.57 | 0.69 | Caecilian |
| 吃皮小蚓螈 | Tiny Cayenne Caecilian | *Microcaecilia unicolor* | Siphonopidae | 4.70 | 1.64 | 0.70 | Caecilian |
| 双线鱼螈 | Two-lined caecilian | *Rhinatrema bivittatum* | Rhinatrematidae | 5.30 | 1.64 | 0.74 | Caecilian |
| 版纳鱼螈 | Banna Caecilian | *Ichthyophis bannanicus* | Ichthyophiidae | 12.20 | 1.45 | 0.67 | Caecilian |

| | | | | | | | |
|---|---|---|---|---|---|---|---|
| 华丽穴居蛙 | Ornate burrowing frog | *Platyplectrum ornatum* | Myobatrachidae | 1.06 | 2.15 | 0.87 | Frogs |
| 热带爪蟾 | western clawed frog | *Xenopus tropicalis* | Pipidae | 1.50 | 2.12 | 0.86 | Frogs |
| 高山倭蛙 | Tibetan frog | *Nanorana parkeri* | Dicroglossidae | 2.30 | 2.14 | 0.86 | Frogs |
| 非洲爪蟾 | African clawed frog | *Xenopus laevis* | Pipidae | 2.70 | 2.06 | 0.85 | Frogs |
| 黑蹼树蛙 | Black-webbed Treefrog | *Rhacophorus kio* | Rhacophoridae | 2.70 | 1.95 | 0.83 | Frogs |
| 宝兴树蛙 | Sichuan whipping frog | *Zhangixalus dugritei* | Rhacophoridae | 3.40 | 2.04 | 0.85 | Frogs |
| 雷山髭蟾 | Leishan moustache toad | *Leptobrachium leishanense* | Megophryidae | 3.50 | 2.00 | 0.84 | Frogs |
| 中华大蟾蜍 | Bufo bufo gargarizans | *Bufo gargarizans* | Bufonidae | 5.40 | 1.96 | 0.82 | Frogs |
| 美洲牛蛙 | American bullfrog | *Lithobates catesbeianus* | Ranidae | 6.30 | 2.09 | 0.86 | Frogs |
| 箭毒蛙 | strawberry poison arrow frog | *Oophaga pumilio* | Dendrobatidae | 6.76 | 1.90 | 0.81 | Frogs |
| 山暗蝾螈 | Allegheny mountain dusky salamader | *Desmognathus ochrophaeus* | Plethodontidae | 15.00 | 1.61 | 0.71 | Salamanders |
| 新疆北鲵 | Central Asian Salamander | ***Ranodon sibiricus*** | Hynobiidae | 21.00 | 1.81 | 0.80 | Salamanders |
| 棕黑疣螈 | Longchuan Crocodile Newt | ***Tylototriton verrucosus*** | Salamandridae | 24.00 | 1.88 | 0.80 | Salamanders |
| 黑腹细长蝾螈 | Black-bellied slender salamander | *Batrachoseps nigriventris* | Plethodontidae | 25.00 | 2.18 | 0.86 | Salamanders |
| 墨西哥钝口螈 | axolotl | *Ambystoma mexicanum* | Ambystomatidae | 32.00 | 1.98 | 0.83 | Salamanders |
| 黑斑肥螈 | Pachytriton | ***Pachytriton brevipes*** | Salamandridae | 40.00 | 2.07 | 0.85 | Salamanders |
| 东方蝾螈 | Qianshan Fire-bellied Newt | ***Cynops orientalis*** | Salamandridae | 44.00 | 2.09 | 0.86 | Salamanders |
| 斑点黑蝾螈 | speckled black salamander | *Aneides flavipunctatus* | Plethodontidae | 44.00 | 1.96 | 0.78 | Salamanders |
| 大鲵 | Chinese giant salamander | ***Andrias davidianus*** | Cryptobranchidae | 48.00 | 1.92 | 0.81 | Salamanders |
| 香港蝾螈 | Hong Kong Warty Newt | ***Paramesotriton hongkongensis*** | Salamandridae | 51.00 | 2.16 | 0.87 | Salamanders |
| 美国隐鳃鲵 | Hellbender | *Cryptobranchus alleganiensis* | Cryptobranchidae | 55.00 | 2.02 | 0.84 | Salamanders |
| 响尾蛇 | prairie rattlesnake | *Crotalus viridis* | Viperidae | 1.30 | 1.98 | 0.82 | Non-avian reptiles |
| 束带蛇 | Coluber sirtalis | *Thamnophis sirtalis* | Colubridae | 1.42 | 2.04 | 0.84 | Non-avian reptiles |
| 蚺 | Burmese python | *Python bivittatus* | Pythonidae | 1.44 | 1.95 | 0.81 | Non-avian reptiles |
| 绿蜥蜴 | green anole | *Anolis carolinensis* | Dactyloidae | 1.70 | 2.13 | 0.86 | Non-avian reptiles |
| 玉米蛇 | Red cornsnake | *Pantherophis guttatus* | Colubridae | 1.73 | 1.99 | 0.82 | Non-avian reptiles |
| 印度眼镜蛇 | India cobra | *Naja naja* | Elapidae | 1.79 | 1.81 | 0.76 | Non-avian reptiles |

| | | | | | | | |
|---|---|---|---|---|---|---|---|
| 温泉蛇 | hot-spring snake | *Thermophis baileyi* | Colubridae | 1.85 | 2.02 | 0.83 | Non-avian reptiles |
| 环形海蛇 | blue-banded sea snake | *Hydrophis cyanocinctus* | Elapidae | 2.02 | 1.81 | 0.76 | Non-avian reptiles |
| 平颏海蛇 | Shaw's sea snake | *Hydrophis curtus* | Elapidae | 2.03 | 2.05 | 0.85 | Non-avian reptiles |
| 扬子鳄 | Chinese alligator | *Alligator sinensis* | Alligatoridae | 2.10 | 1.84 | 0.80 | Non-avian reptiles |
| 鳄鱼 | Siamese Crocodile | *Crocodylus siamensis* | Crocodylinae | 2.10 | 1.81 | 0.79 | Non-avian reptiles |
| 乌龟 | three-keeled pond turtle | *Mauremys reevesii* | Geoemydidae | 2.20 | 1.95 | 0.82 | Non-avian reptiles |
| 鳄蜥 | crocodile lizard | *Shinisaurus crocodilurus* | Shinisauridae | 2.20 | 1.79 | 0.76 | Non-avian reptiles |
| 悬崖壁蜥 | Cape cliff lizard | *Hemicordylus capensis* | Cordylidae | 2.30 | 1.73 | 0.75 | Non-avian reptiles |
| 三趾箱龟 | Three-toed box turtle | *Terrapene carolina* | Emydidae | 4.18 | 1.96 | 0.83 | Non-avian reptiles |
| 大蜥蜴 | tuatara | *Sphenodon punctatus* | Sphenodontidae | 5.00 | 1.69 | 0.71 | Non-avian reptiles |
| 环颈雉 | Ring-necked pheasant | *Phasianus colchicus* | Phasianidae | 0.90 | 0.89 | 0.43 | Birds |
| 鸡 | chicken | *Gallus gallus* | Phasianidae | 1.10 | 0.89 | 0.45 | Birds |
| 鸭 | duck | *Anas anas* | Anatidae | 1.10 | 0.60 | 0.26 | Birds |
| 双领鸻 | killdeer | *Charadrius vociferus* | Charadriidae | 1.20 | 0.62 | 0.33 | Birds |
| 绒毛啄木鸟 | Downy woodpecker | *Dryobates pubescens* | Picidae | 1.20 | 0.36 | 0.16 | Birds |
| 白尾鹲 | White-tailed Tropicbird | *Phaethon lepturus* | Phaethontidae | 1.20 | 0.73 | 0.42 | Birds |
| 红冠鹦鹉 | Red-crested Turaco | *Tauraco erythrolophus* | Musophagidae | 1.20 | 0.72 | 0.40 | Birds |
| 斑马雀 | zebra finch | *Taeniopygia guttata* | Estrildidae | 1.20 | 0.84 | 0.53 | Birds |
| 纹腹鹰 | Sharp-shinned hawk | *Accipiter striatus* | Accipitridae | 1.55 | 1.03 | 0.57 | Birds |
| 美洲角雕 | harpy eagle | *Harpia harpyja* | Accipitridae | 1.58 | 1.01 | 0.44 | Birds |
| 亚马逊白头鹦鹉 | Cuban parrot | *Amazona leucocephala* | Psittacidae | 1.65 | 0.75 | 0.58 | Birds |
| 小凤头鹦鹉 | Little corella | *Cacatua sanguinea* | Psittacidae | 1.71 | 0.74 | 0.39 | Birds |
| 欧亚雕鸮 | Eurasian eagle owl | *Bubo bubo* | Strigidae | 2.03 | 0.90 | 0.48 | Birds |
| 鸵鸟 | South African ostrich | *Struthio australis* | Struthionidae | 2.16 | 0.75 | 0.28 | Birds |
| 鸭嘴兽 | platypus | *Ornithorhynchus anatinus* | Ornithorhynchidae | 1.99 | 0.96 | 0.55 | Mammals |
| 针鼹 | short-beaked echidna | *Tachyglossus aculeatus* | Tachyglossidae | 2.20 | 0.96 | 0.55 | Mammals |
| 大熊猫 | giant panda | *Ailuropoda melanoleuca* | Ursidae | 2.20 | 1.56 | 0.74 | Mammals |

| | | | | | | | |
|---|---|---|---|---|---|---|---|
| 大狐蝠 | large flying fox | *Pteropus vampyrus* | Pteropodidae | 2.20 | 1.44 | 0.64 | Mammals |
| 狗 | dogs | *Canis lupus familiaris* | Canidae | 2.39 | 1.15 | 0.56 | Mammals |
| 猪 | pig | *Sus scrofa* | Suidae | 2.50 | 1.44 | 0.70 | Mammals |
| 小袋鼠 | Quokka | *Setonix brachyurus* | Macropodidae | 2.50 | 1.58 | 0.76 | Mammals |
| 小鼠 | house mouse | *Mus musculus* | Muridae | 2.70 | 1.24 | 0.65 | Mammals |
| 鲸 | northern bottlenose whale | *Hyperoodon ampullatus* | Ziphiidae | 2.80 | 1.01 | 0.45 | Mammals |
| 人 | human | *Homo sapiens* | Hominidae | 3.00 | 1.68 | 0.77 | Mammals |
| 考拉 | Koala | *Phascolarctos cinereus* | Phascolarctidae | 3.20 | 1.55 | 0.72 | Mammals |
| 亚洲象 | Asian elephant | *Elephas maximus* | Elephantidae | 3.20 | 1.52 | 0.72 | Mammals |
| 象鼩 | Cape elephant shrew | *Elephantulus edwardii* | Macroscelididae | 4.20 | 1.15 | 0.58 | Mammals |
| 金毛鼹 | Cape golden mole | *Chrysochloris asiatica* | Chrysochloridae | 4.70 | 1.37 | 0.71 | Mammals |

#### S6 Relationship between the abundance and expression of TE superfamilies

| Species | Relationship between abundance (bp, x) of TE superfamilies in the genome and their expression (y) in the gonads | | | | | |
|---|---|---|---|---|---|---|
| | Equation of relationship | | Comparison of correlation between sex | | | |
| | Testis | Ovary | *t* (Slope) | *P* (Slope) | *t* (Intercept) | *P* (Intercept) |
| RS | log(y) = **0.6555** log(x) -2.2045 | log(y) =0.7215 log(x) **-3.0247** | 0.1227 | 0.9032 | -0.8655 | 0.3941 |
| TV | log(y) = 0.9734 log(x) -4.2162 | log(y) =1.0496 log(x) -5.4483 | 0.2029 | 0.8409 | -2.2149 | 0.0361 |
| PB | log(y) = 1.0671 log(x) -5.4918 | log(y) =0.9395 log(x) -5.0998 | -0.2754 | 0.7850 | -1.7232 | 0.0955 |
| CO | log(y) = **1.3530** log(x) -8.1203 | log(y) =1.0665 log(x) -6.3788 | -0.6398 | 0.5271 | -1.3564 | 0.1851 |
| AD | log(y) = 0.7841 log(x) -3.0878 | log(y) =0.8658 log(x) -4.1511 | 0.4360 | 0.6664 | -2.4164 | 0.0230 |
| PH | log(y) =1.1394 log(x) -5.9990 | log(y) =**1.1321** log(x) -6.3466 | -0.0170 | 0.9866 | -1.2679 | 0.2135 |

| Analysis of covariance | Comparison of slopes for males across species | | Comparison of slopes for females across species | |
|---|---|---|---|---|
| | Heterogeneity of slopes | intercept | Heterogeneity of slopes | intercept |
| *F* | 0.739 | 28.822 | 0.284 | 37.323 |
| *P* | 0.596 | **< 0.001** | 0.921 | **< 0.001** |

**S7 The 10 most highly expressed TE superfamilies in the six salamander species**

| species | Top 10 expressed superfamilies in females | Average expression in Females (TPM) | Top 10 expressed superfamilies in males | Average expression in males (TPM) |
|---|---|---|---|---|
| *R. sibiricus* | LINE/Jockey | 6206 | LINE/Jockey | 15535 |
| *R. sibiricus* | DIRS/DIRS | 5427 | DIRS/DIRS | 11962 |
| *R. sibiricus* | LINE/L1 | 2954 | LINE/L1 | 7610 |
| *R. sibiricus* | LTR/Gypsy | 827 | LTR/Gypsy | 3024 |
| *R. sibiricus* | TIR/Harbinger | 575 | TIR/Harbinger | 1359 |
| *R. sibiricus* | LTR/ERV | 392 | Maverick/Maverick | 762 |
| *R. sibiricus* | LINE/RTE | 130 | LTR/ERV | 694 |
| *R. sibiricus* | PLE/Penelope | 123 | LINE/RTE | 360 |
| *R. sibiricus* | TIR/hAT | 83 | PLE/Penelope | 352 |
| *R. sibiricus* | TIR/Mariner | 58 | LINE/I | 125 |
| *T. verrucosus* | LTR/Gypsy | 1495 | DIRS/DIRS | 12179 |
| *T. verrucosus* | DIRS/DIRS | 1381 | LTR/Gypsy | 6162 |
| *T. verrucosus* | LINE/L1 | 1302 | LINE/L1 | 5666 |
| *T. verrucosus* | LINE/Jockey | 759 | LINE/Jockey | 4104 |
| *T. verrucosus* | TIR/Harbinger | 524 | LTR/ERV | 2870 |
| *T. verrucosus* | TIR/hAT | 372 | TIR/Harbinger | 1635 |
| *T. verrucosus* | LTR/ERV | 314 | PLE/Penelope | 901 |
| *T. verrucosus* | PLE/Penelope | 270 | LINE/RTE | 881 |
| *T. verrucosus* | LINE/RTE | 184 | Maverick/Maverick | 561 |
| *T. verrucosus* | LTR/Retrovirus | 101 | LTR/Retrovirus | 408 |
| *P. brevipes* | LINE/L1 | 869 | DIRS/DIRS | 10115 |
| *P. brevipes* | DIRS/DIRS | 852 | LINE/L1 | 4021 |

| | | | | |
|---|---|---|---|---|
| *P. brevipes* | LTR/Gypsy | 828 | LINE/Jockey | 3928 |
| *P. brevipes* | TIR/hAT | 742 | LTR/Gypsy | 3595 |
| *P. brevipes* | LINE/Jockey | 634 | LTR/ERV | 2773 |
| *P. brevipes* | LTR/ERV | 445 | LTR/Retrovirus | 464 |
| *P. brevipes* | TIR/Harbinger | 386 | PLE/Penelope | 412 |
| *P. brevipes* | LINE/RTE | 108 | TIR/hAT | 385 |
| *P. brevipes* | PLE/Penelope | 94 | LINE/RTE | 327 |
| *P. brevipes* | TIR/PiggyBac | 60 | TIR/Harbinger | 293 |
| *C. orientalis* | LINE/L1 | 1522 | DIRS/DIRS | 9592 |
| *C. orientalis* | LTR/Gypsy | 1211 | LINE/L1 | 7674 |
| *C. orientalis* | DIRS/DIRS | 1085 | LINE/Jockey | 5033 |
| *C. orientalis* | LINE/Jockey | 854 | LTR/Gypsy | 4245 |
| *C. orientalis* | TIR/hAT | 399 | LTR/ERV | 1649 |
| *C. orientalis* | PLE/Penelope | 357 | PLE/Penelope | 1143 |
| *C. orientalis* | LINE/RTE | 217 | LINE/RTE | 692 |
| *C. orientalis* | LTR/ERV | 167 | LTR/Retrovirus | 221 |
| *C. orientalis* | TIR/Harbinger | 83 | TIR/Harbinger | 209 |
| *C. orientalis* | TIR/PiggyBac | 68 | TIR/hAT | 180 |
| *A. davidianus* | DIRS/DIRS | 14429 | DIRS/DIRS | 17415 |
| *A. davidianus* | LINE/Jockey | 3960 | LINE/Jockey | 6722 |
| *A. davidianus* | LTR/Gypsy | 1052 | LINE/L1 | 4497 |
| *A. davidianus* | LINE/L1 | 1012 | LTR/Gypsy | 4169 |
| *A. davidianus* | PLE/Penelope | 615 | Helitron/Helitron | 1223 |
| *A. davidianus* | TIR/hAT | 157 | PLE/Penelope | 1094 |
| *A. davidianus* | Maverick/Maverick | 117 | LTR/ERV | 493 |
| *A. davidianus* | TIR/Mariner | 106 | TIR/Harbinger | 177 |

| | | | | |
|---|---|---|---|---|
| *A. davidianus* | LTR/ERV | 102 | TIR/hAT | 156 |
| *A. davidianus* | TIR/Harbinger | 80 | Maverick/Maverick | 147 |
| *P. hongkongensis* | DIRS/DIRS | 5526 | DIRS/DIRS | 20239 |
| *P. hongkongensis* | LINE/L1 | 3737 | LINE/L1 | 13187 |
| *P. hongkongensis* | LINE/Jockey | 2412 | LINE/Jockey | 7272 |
| *P. hongkongensis* | LTR/Gypsy | 2105 | LTR/Gypsy | 6415 |
| *P. hongkongensis* | LTR/ERV | 1023 | LTR/ERV | 3795 |
| *P. hongkongensis* | PLE/Penelope | 664 | PLE/Penelope | 2254 |
| *P. hongkongensis* | TIR/hAT | 423 | LINE/RTE | 1220 |
| *P. hongkongensis* | LINE/RTE | 393 | LTR/Retrovirus | 784 |
| *P. hongkongensis* | LTR/Retrovirus | 249 | TIR/Harbinger | 386 |
| *P. hongkongensis* | TIR/Harbinger | 192 | Maverick/Maverick | 274 |

**S8 Overall expression levels of genes and TEs in the ovary or testis of the six salamander species (% TPM)**

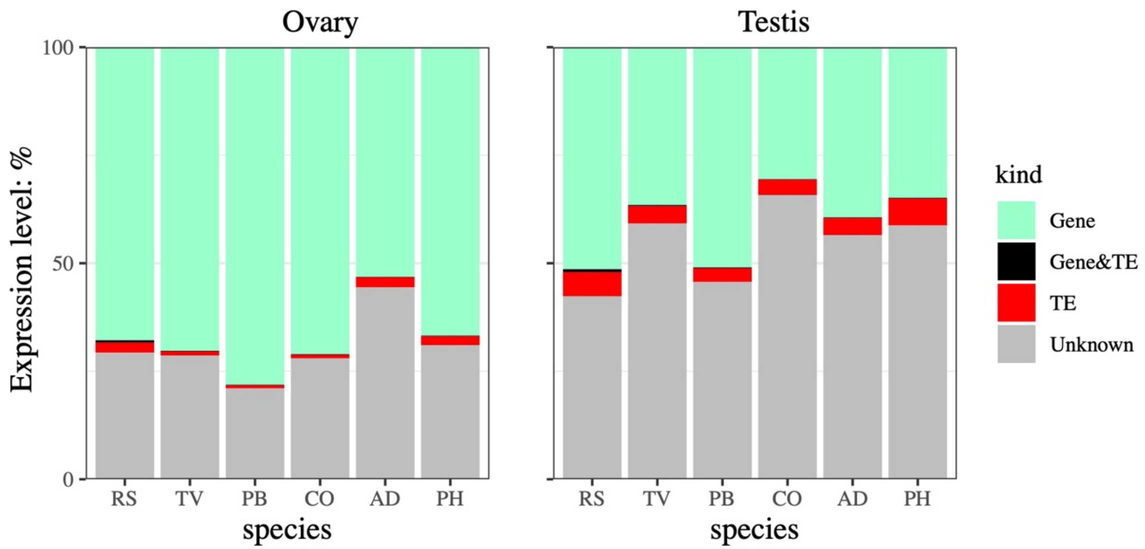

Species are arranged left to right by increasing genome size: RS, *Ranodon sibiricus*; TV, *Tylototriton verrucosus*; PB, *Pachytriton brevipes*; CO, *Cynops orientalis*; AD, *Andrias davidianus*; PH, *Paramesotriton hongkongensis*

**S9 Length distribution (A) and 5' U bias (B) of small RNA molecules in the gonads of individual male and female salamander samples.**

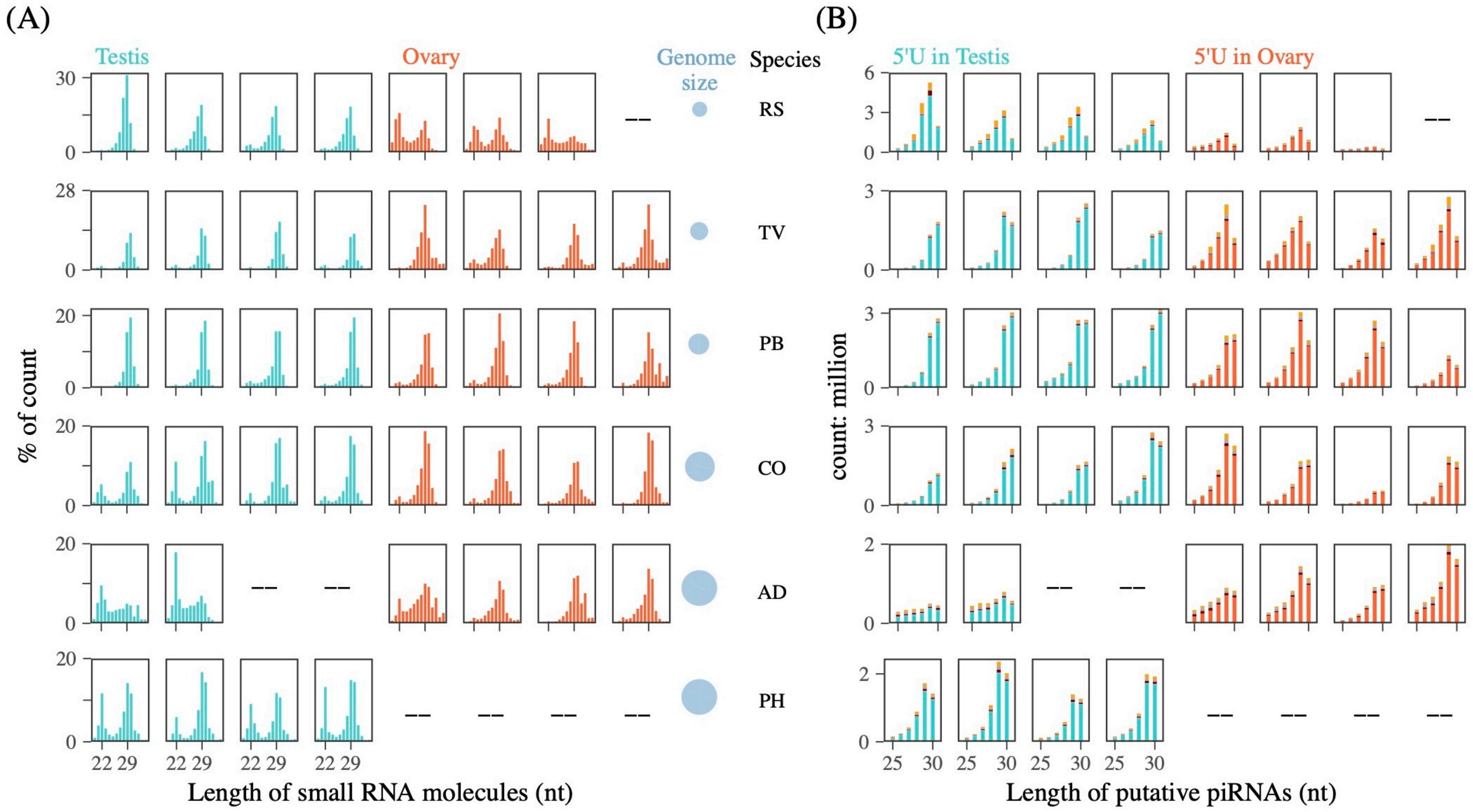

#### S10 The clean small RNAs and putative piRNAs in each salamander sample

| species | sex | sample | clean_18-40 nt | Total putative piRNAs (25-30 nt) | Unique putative piRNA (25-30 nt) | Total piRNA (25-30)/total clean (18-40 nt) | Average Total piRNA (25-30 nt) / total clean (18-40 nt) | Unique piRNA /total piRNAs (25-30 nt) | **Normalized uniq piRNA (per 10M small RNAs 18-40 nt in length)** | **Normalized unique piRNAs mapping to TEs** |
|---|---|---|---|---|---|---|---|---|---|---|
| *Ranodon sibiricus* | female | WQ02 | 11,604,083 | 4,868,129 | 496,496 | 42.0% | 37% | 10.2% | 427,863 | 97,510 |
| | female | WQ03 | 13,591,993 | 5,581,827 | 779,671 | 41.1% | | 14.0% | 573,625 | 154,936 |
| | female | WQ04 | 6,896,722 | 2,003,438 | 266,851 | 29.0% | | 13.3% | 386,924 | 71,349 |
| | male | WQ05 | 16,935,126 | 13,288,223 | 470,649 | 78.5% | 60% | 3.5% | 277,913 | 42,826 |
| | male | WQ07 | 16,496,516 | 9,374,629 | 986,996 | 56.8% | | 10.5% | 598,306 | 140,662 |
| | male | WQ08 | 18,341,802 | 9,814,338 | 905,599 | 53.5% | | 9.2% | 493,735 | 101,660 |
| | male | WQ10 | 13,082,664 | 6,940,705 | 774,833 | 53.1% | | 11.2% | 592,259 | 138,529 |
| *Tylototriton verrucosus* | female | HS08 | 10,838,684 | 6,590,103 | 395,443 | 60.8% | 53% | 6.0% | 364,844 | 98,070 |
| | female | HS09 | 14,332,777 | 6,818,958 | 904,658 | 47.6% | | 13.3% | 631,181 | 206,017 |
| | female | HS11 | 9,689,432 | 4,312,095 | 496,054 | 44.5% | | 11.5% | 511,954 | 159,218 |
| | female | HS12 | 12,501,636 | 7,527,884 | 417,748 | 60.2% | | 5.5% | 334,155 | 92,093 |
| | male | HS02 | 13,921,523 | 3,858,594 | 839,636 | 27.7% | 33% | 21.8% | 603,121 | 176,292 |
| | male | HS04 | 15,021,365 | 5,336,718 | 1,091,850 | 35.5% | | 20.5% | 726,865 | 217,841 |
| | male | HS05 | 14,815,435 | 5,445,816 | 1,007,806 | 36.8% | | 18.5% | 680,241 | 211,419 |
| | male | HS07 | 11,682,213 | 3,650,348 | 756,193 | 31.2% | | 20.7% | 647,303 | 198,528 |
| *Pachytriton brevipes* | female | NF08 | 10,245,368 | 3,469,373 | 354,227 | 33.9% | 46% | 10.2% | 345,744 | 47,263 |
| | female | NF09 | 14,267,591 | 6,234,115 | 921,343 | 43.7% | | 14.8% | 645,759 | 89,890 |
| | female | NF20 | 14,800,637 | 8,157,582 | 1,015,889 | 55.1% | | 12.5% | 686,382 | 98,564 |
| | female | NF21 | 14,697,533 | 7,341,309 | 974,122 | 49.9% | | 13.3% | 662,779 | 97,362 |
| | male | NF01 | 14,346,946 | 6,047,212 | 1,020,091 | 42.1% | 44% | 16.9% | 711,016 | 117,460 |

| | | | | | | | | | | |
|---|---|---|---|---|---|---|---|---|---|---|
| | male | NF14 | 16,313,396 | 7,178,965 | 1,166,880 | 44.0% | | 16.3% | 715,289 | 116,807 |
| | male | NF16 | 17,477,879 | 7,790,731 | 1,241,258 | 44.6% | | 15.9% | 710,188 | 113,914 |
| | male | NF18 | 16,367,997 | 7,599,481 | 1,142,246 | 46.4% | | 15.0% | 697,853 | 108,656 |
| *Cynops orientalis* | female | CO12 | 14,482,941 | 7,682,291 | 951,754 | 53.0% | 44% | 12.4% | 657,155 | 123,282 |
| | female | CO13 | 12,185,835 | 5,083,594 | 723,200 | 41.7% | | 14.2% | 593,476 | 93,710 |
| | female | CO14 | 5,146,586 | 1,704,233 | 293,067 | 33.1% | | 17.2% | 569,440 | 84,505 |
| | female | QZ09 | 10,100,463 | 4,939,582 | 684,507 | 48.9% | | 13.9% | 677,699 | 103,417 |
| | male | CO16 | 11,142,943 | 2,926,582 | 561,094 | 26.3% | 39% | 19.2% | 503,542 | 72,258 |
| | male | CO17 | 13,176,247 | 4,998,264 | 670,222 | 37.9% | | 13.4% | 508,659 | 80,877 |
| | male | QZ01 | 9,725,994 | 4,175,352 | 589,702 | 42.9% | | 14.1% | 606,315 | 116,716 |
| | male | QZ04 | 15,768,386 | 7,463,655 | 1,057,250 | 47.3% | | 14.2% | 670,487 | 123,101 |
| *Andrias davidianus* | female | LY01 | 9,035,740 | 3,683,819 | 730,110 | 40.8% | 38% | 19.8% | 808,025 | 161,605 |
| | female | LY03 | 13,542,759 | 4,628,997 | 782,411 | 34.2% | | 16.9% | 577,734 | 157,548 |
| | female | LY04 | 8,088,094 | 2,798,328 | 487,252 | 34.6% | | 17.4% | 602,431 | 162,355 |
| | female | LY05 | 14,577,753 | 6,223,416 | 969,496 | 42.7% | | 15.6% | 665,052 | 185,283 |
| | male | LY06 | 7,339,454 | 1,018,559 | 187,174 | 13.9% | 18% | 18.4% | 255,024 | 51,107 |
| | male | LY07 | 6,929,866 | 240,388 | 48,312 | 3.5% | | 20.1% | 69,716 | 12,786 |
| | male | LY08 | 10,126,404 | 2,313,631 | 421,496 | 22.8% | | 18.2% | 416,235 | 90,032 |
| | male | LY09 | 11,471,238 | 3,440,088 | 829,412 | 30.0% | | 24.1% | 723,036 | 148,005 |
| *Paramesotriton honkongensis* | male | HK03 | 12,200,956 | 4,848,719 | 671,892 | 39.7% | 39% | 13.9% | 550,688 | 109,477 |
| | male | HK04 | 14,084,143 | 6,242,456 | 894,350 | 44.3% | | 14.3% | 635,005 | 136,018 |
| | male | HK05 | 11,872,338 | 3,769,704 | 614,821 | 31.8% | | 16.3% | 517,860 | 103,468 |
| | male | HK06 | 13,394,295 | 5,533,221 | 687,128 | 41.3% | | 12.4% | 513,000 | 98,650 |

**S11 Very few unique piRNAs are shared between species**

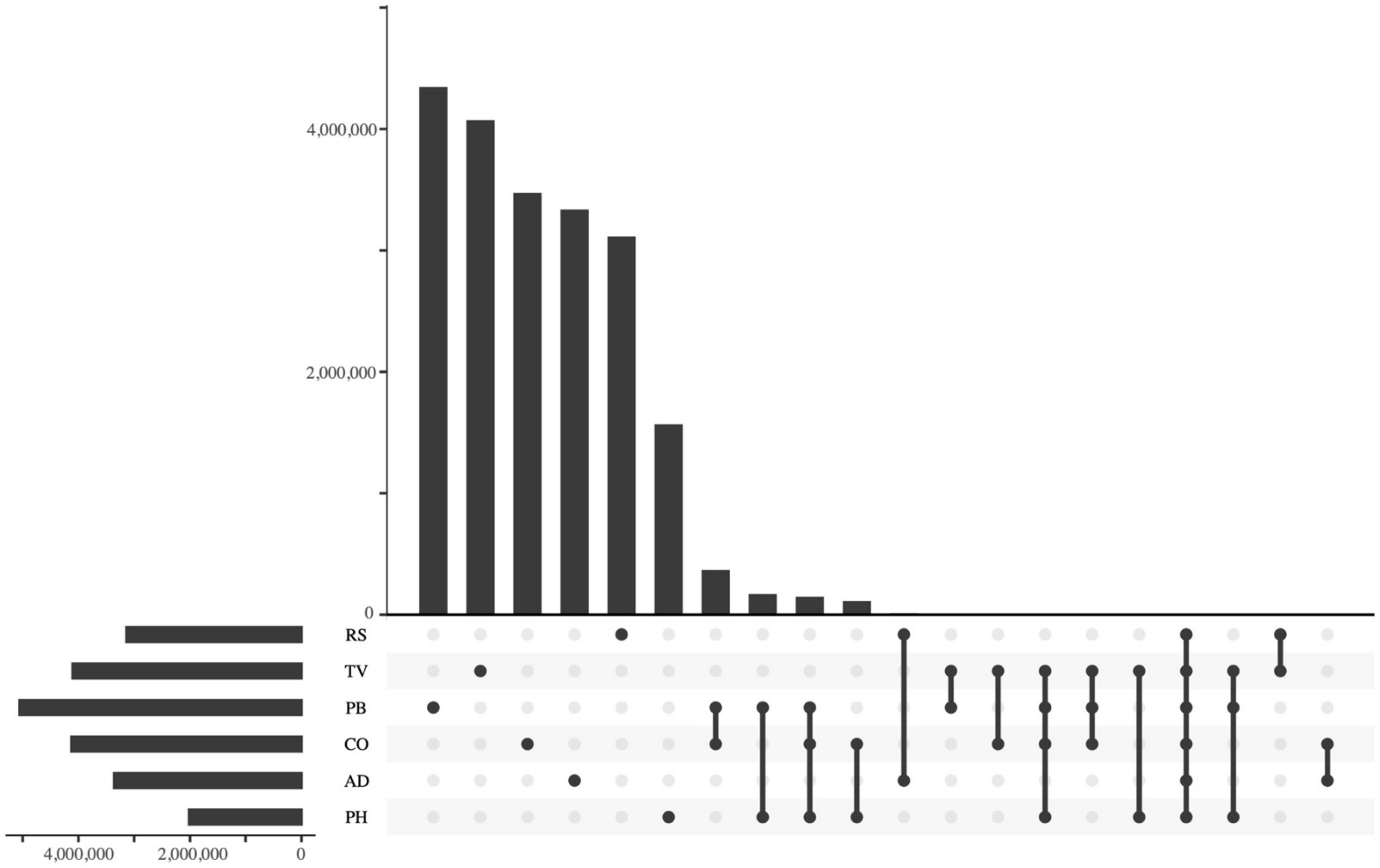

Species are arranged top to bottom by increasing genome size: RS, *Ranodon sibiricus*; TV, *Tylototriton verrucosus*; PB, *Pachytriton brevipes*; CO, *Cynops orientalis*; AD, *Andrias davidianus*; PH, *Paramesotriton hongkongensis.* Vertical bars show the number of unique piRNAs in each species (columns 1-6, with a black dot indicating the species) or shared between two or more species (black dots and lines connecting them). Columns 11-20 include fewer than 111, 314 unique piRNA sequences.

#### S12 Relationship between the expression of TE superfamilies and piRNAs mapping to TE superfamilies

| Species | Relationship between expression (TPM, x) of TE superfamilies and the piRNAs mapped to TE superfamilies (y) | | | | | |
|---|---|---|---|---|---|---|
| | Equation of relationship | | Comparison of correlation between sex | | | |
| | Testis | Ovary | *t* (Slope) | *P* (Slope) | *t* (Intercept) | *P* (Intercept) |
| RS | log(y) = 0.6235 log(x) + 2.4754 | log(y) = 0.6508 log(x) + 2.4353 | 0.1015 | 0.9197 | 0.0872 | 0.9310 |
| TV | log(y) = 0.6241 log(x) + 2.7548 | log(y) = 0.6296 log(x) + 2.7359 | 0.0244 | 0.9806 | -0.0204 | 0.9838 |
| PB | log(y) = 0.8638 log(x) + 1.9357 | log(y) = 0.5894 log(x) + 3.0540 | -1.1419 | 0.2590 | 1.6104 | 0.1137 |
| CO | log(y) = **0.9118** log(x) + 1.8017 | log(y) = 0.5867 log(x) + 3.0809 | -1.6060 | 0.1144 | 2.2127 | 0.0314 |
| AD | log(y) = 0.8454 log(x) + 1.9993 | log(y) = **0.8853** log(x) + 1.8208 | 0.1749 | 0.8620 | -0.3010 | 0.7649 |
| PH | log(y) = 0.8939 log(x) + 1.5568 | | | | | |

| Analysis of covariance | Comparison of slopes for males across species | | Comparison of slopes for females across species | |
|---|---|---|---|---|
| | Heterogeneity of slopes | intercept | Heterogeneity of slopes | intercept |
| *F* | 0.818 | 168.825 | 0.356 | 153.621 |
| *P* | 0.539 | **< 0.001** | 0.84 | **< 0.001** |

**S13 The average expression levels of all putative piRNA clusters in each salamander species.**

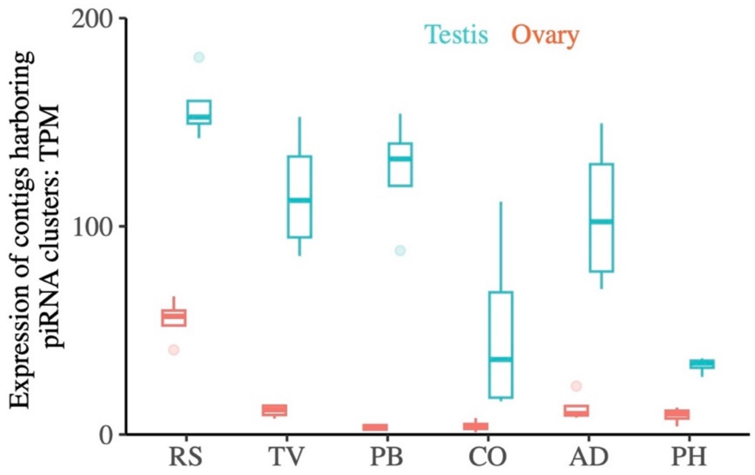

Species are arranged left to right by increasing genome size: RS, *Ranodon sibiricus*; TV, *Tylototriton verrucosus*; PB, *Pachytriton brivipes*; CO, *Cynops orientalis*; AD, *Andrias davidianus*; PH, *Paramesotriton hongkongensis*.

**S14 Results of the *de novo* transcriptome assembly.**

| Species | Number of transcripts | Sum length | Min length | Average length | Max length | N50 of transcripts |
|---|---|---|---|---|---|---|
| *Ranodon sibiricus* | 573,681 | 473,363,550 | 183 | 825 | 28,590 | 1,874 |
| *Tylototriton verrucosus* | 614,818 | 544,285,339 | 175 | 885 | 24,708 | 2,205 |
| *Pachytriton brevipes* | 597,551 | 531,263,346 | 180 | 889 | 39,116 | 2,099 |
| *Cynops orientalis* | 598,805 | 514,032,901 | 182 | 858 | 25,106 | 2,093 |
| *Andrias davidianus* | 452,395 | 401,061,354 | 174 | 887 | 27,377 | 2,099 |
| *Paramesotriton hongkongensis* | 690,114 | 531,632,536 | 168 | 770 | 28,364 | 1,927 |

#### S15 The number of retained small RNA molecules after each processing step.

| Species | Tissue | Samples | RIN value of RNA | Clean reads (after FastX) | After adapter cut (18-40 nt) | riboRNA fragments removed | tRNA fragments removed | clean_18-40 nt | Putative piRNAs (25-30 nt) | piRNAs mapped to TEs (25-30 nt) | Unique piRNAs (25-30 nt) | Normalized uniq piRNA (10M, 18-40) | % unique piRNA mapped to TE transcript | ping pong occurrence (10 bp overlap) |
|---|---|---|---|---|---|---|---|---|---|---|---|---|---|---|
| *Ranodon sibiricus* | Testis#1 | WQ07 | 9.6 | 18,628,420 | 18,387,280 | 1,393,269 | 497,495 | 16,496,516 | 9,374,629 | 1,621,347 | 986,996 | 598,306 | 23.51% | 11,766 |
| | Testis#2 | WQ08 | 8.9 | 20,632,085 | 20,127,121 | 702,889 | 1,082,430 | 18,341,802 | 9,814,338 | 1,111,036 | 905,599 | 493,735 | 20.59% | 8,230 |
| | Testis#3 | WQ10 | 9.4 | 15,341,619 | 15,148,223 | 1,731,806 | 333,753 | 13,082,664 | 6,940,705 | 1,189,737 | 774,833 | 592,259 | 23.39% | 7,520 |
| | Testis#4 | WQ05 | 9.3 | 18,141,749 | 18,022,557 | 682,282 | 405,149 | 16935126 | 13,288,223 | 582,031 | 470,649 | 277,913 | 15.41% | 3,398 |
| | Ovary#1 | WQ03 | 9.3 | 17,133,153 | 16,652,089 | 2,708,832 | 351,264 | 13,591,993 | 5,581,827 | 1,835,311 | 779,671 | 573,625 | 27.01% | 10,287 |
| | Ovary#2 | WQ02 | 9.6 | 15,112,304 | 14,382,824 | 2,298,340 | 480,401 | 11604083 | 4,868,129 | 1,518,581 | 496,496 | 427,863 | 22.79% | 2,735 |
| | Ovary#3 | WQ04 | 8.7 | 13,323,488 | 11,664,671 | 4,260,265 | 507,684 | 6896722 | 2,003,438 | 810,078 | 266,851 | 386,924 | 18.44% | 1,678 |
| *Tylototriton verrucosus* | Testis#1 | HS02 | 9.3 | 15,812,676 | 15,593,182 | 551,836 | 1,119,823 | 13,921,523 | 3,858,594 | 1,093,581 | 839,636 | 603,121 | 29.23% | 4,047 |
| | Testis#2 | HS04 | 9.2 | 16,818,923 | 16,603,246 | 475,376 | 1,106,505 | 15,021,365 | 5,336,718 | 1,558,405 | 1,091,850 | 726,865 | 29.97% | 6,056 |
| | Testis#3 | HS05 | 9.6 | 16,893,170 | 16,721,870 | 277,675 | 1,628,760 | 14,815,435 | 5,445,816 | 1,688,611 | 1,007,806 | 680,241 | 31.08% | 5,942 |
| | Testis#4 | HS07 | 9.3 | 15,502,396 | 15,246,506 | 445,413 | 3,118,880 | 11,682,213 | 3,650,348 | 1,088,187 | 756,193 | 647,303 | 30.67% | 3,803 |
| | Ovary#1 | HS08 | 6.5 | 30,369,861 | 15,511,083 | 3,212,333 | 1,460,066 | 10,838,684 | 6,590,103 | 2,272,313 | 395,443 | 364,844 | 26.88% | 2,216 |
| | Ovary#2 | HS09 | 8.5 | 17,236,506 | 16,690,699 | 1,520,090 | 837,832 | 14,332,777 | 6,818,958 | 2,553,943 | 904,658 | 631,181 | 32.64% | 10,485 |
| | Ovary#3 | HS11 | 7.8 | 16,009,250 | 14,887,546 | 2,502,646 | 2,695,468 | 9,689,432 | 4,312,095 | 1,458,617 | 496,054 | 511,954 | 31.10% | 3,859 |
| | Ovary#4 | HS12 | 7.8 | 34,354,037 | 16,987,985 | 3,450,516 | 1,035,833 | 12,501,636 | 7,527,884 | 2,725,514 | 417,748 | 334,155 | 27.56% | 2,187 |
| *Pachytriton brevipes* | Testis#1 | NF01 | 9.4 | 16,480,564 | 16,288,267 | 420,115 | 1,521,206 | 14,346,946 | 6,047,212 | 975,057 | 1,020,091 | 711,016 | 16.52% | 3,043 |
| | Testis#2 | NF14 | 9.8 | 17,411,312 | 17,244,483 | 538,568 | 392,519 | 16,313,396 | 7,178,965 | 1,151,729 | 1,166,880 | 715,289 | 16.33% | 3,829 |
| | Testis#3 | NF16 | 9.7 | 18,801,974 | 18,559,761 | 727,164 | 354,718 | 17,477,879 | 7,790,731 | 1,198,551 | 1,241,258 | 710,188 | 16.04% | 4,015 |
| | Testis#4 | NF18 | 9.7 | 17,318,113 | 17,116,842 | 345,509 | 403,336 | 16,367,997 | 7,599,481 | 1,070,642 | 1,142,246 | 697,853 | 15.57% | 3,320 |
| | Ovary#1 | NF08 | 8.5 | 31,360,428 | 12,576,185 | 99,238 | 2,231,579 | 10,245,368 | 3,469,373 | 497,323 | 354,227 | 345,744 | 13.67% | 689 |
| | Ovary#2 | NF09 | 8.3 | 16,735,557 | 16,200,399 | 1,187,120 | 745,688 | 14,267,591 | 6,234,115 | 890,850 | 921,343 | 645,759 | 13.92% | 3,934 |
| | Ovary#3 | NF20 | 8.7 | 16,919,838 | 16,544,102 | 1,032,781 | 710,684 | 14,800,637 | 8,157,582 | 1,216,667 | 1,015,889 | 686,382 | 14.36% | 4,844 |
| | Ovary#4 | NF21 | 8.3 | 17,317,850 | 16,958,860 | 1,749,003 | 512,324 | 14,697,533 | 7,341,309 | 1,123,775 | 974,122 | 662,779 | 14.69% | 4,227 |

| | | | | | | | | | | | | | | |
|---|---|---|---|---|---|---|---|---|---|---|---|---|---|---|
| *Cynops orientali* | Testis#1 | CO16 | 8.3 | 16,241,320 | 15,527,596 | 667,498 | 3,717,155 | 11,142,943 | 2,926,582 | 381,623 | 561,094 | 503,542 | 14.35% | 630 |
| | Testis#2 | CO17 | 8.1 | 18,577,955 | 16,985,684 | 1,780,592 | 2,028,845 | 13,176,247 | 4,998,264 | 830,554 | 670,222 | 508,659 | 15.90% | 1,525 |
| | Testis#3 | QZ01 | 7.6 | 14,530,233 | 14,401,574 | 555,596 | 4,119,984 | 9,725,994 | 4,175,352 | 786,848 | 589,702 | 606,315 | 19.25% | 1,706 |
| | Testis#4 | QZ04 | 8.1 | 17,831,129 | 17,552,203 | 880,748 | 903,069 | 15,768,386 | 7,463,655 | 1,405,890 | 1,057,250 | 670,487 | 18.36% | 3,669 |
| | Ovary#1 | CO12 | 8.5 | 15,716,060 | 15,427,646 | 279,633 | 665,072 | 14,482,941 | 7,682,291 | 1,500,478 | 951,754 | 657,155 | 18.76% | 5,606 |
| | Ovary#2 | CO13 | 6.7 | 17,546,503 | 17,220,001 | 251,580 | 4,782,586 | 12,185,835 | 5,083,594 | 717,686 | 723,200 | 593,476 | 15.79% | 2,732 |
| | Ovary#3 | CO14 | 7.4 | 13,607,087 | 13,337,094 | 350,564 | 7,839,944 | 5,146,586 | 1,704,233 | 222,006 | 293,067 | 569,440 | 14.84% | 544 |
| | Ovary#4 | QZ09 | 8.3 | 15,910,436 | 14,940,315 | 1,567,766 | 3,272,086 | 10,100,463 | 4,939,582 | 696,729 | 684,507 | 677,699 | 15.26% | 2,214 |
| *Andrias davidianus* | Testis#1 | LY06 | 8.6 | 15,210,326 | 13,637,209 | 2,595,045 | 3,702,710 | 7,339,454 | 1,018,559 | 126,220 | 187,174 | 255,024 | 20.04% | 723 |
| | Testis#2 | LY07 | 8.2 | 16,624,734 | 11,166,742 | 3,533,085 | 703,791 | 6,929,866 | 240,388 | 27,777 | 48,312 | 69,716 | 18.34% | 96 |
| | Testis#3 | LY08 | 7.2 | 18,033,692 | 16,524,625 | 4,351,823 | 2,046,398 | 10,126,404 | 2,313,631 | 352,947 | 421,496 | 416,235 | 21.63% | 2,609 |
| | Testis#4 | LY09 | 8.7 | 19,154,630 | 17,722,859 | 4,327,063 | 1,924,558 | 11,471,238 | 3,440,088 | 661,005 | 829,412 | 723,036 | 20.47% | 4,707 |
| | Ovary#1 | LY01 | 8.2 | 17,702,705 | 14,359,441 | 3,357,714 | 1,965,987 | 9,035,740 | 3,683,819 | 739,659 | 730,110 | 808,025 | 20.00% | 4,317 |
| | Ovary#2 | LY03 | 7.0 | 17,702,705 | 17,258,351 | 2,589,766 | 1,125,826 | 13,542,759 | 4,628,997 | 1,150,861 | 782,411 | 577,734 | 27.27% | 7,168 |
| | Ovary#3 | LY04 | 7.5 | 17,702,705 | 14,713,470 | 1,477,203 | 5,148,173 | 8,088,094 | 2,798,328 | 722,970 | 487,252 | 602,431 | 26.95% | 3,047 |
| | Ovary#4 | LY05 | 7.3 | 17,702,705 | 17,364,607 | 1,788,881 | 997,973 | 14,577,753 | 6,223,416 | 1,671,421 | 969,496 | 665,052 | 27.86% | 10,004 |
| *Paramesotriton honkongensis* | Testis#1 | HK03 | 9.0 | 15,551,891 | 14,837,493 | 1,095,238 | 1,541,299 | 12,200,956 | 4,848,719 | 902,050 | 671,892 | 550,688 | 19.88% | 1,628 |
| | Testis#2 | HK04 | 8.3 | 17,610,098 | 17,122,533 | 1,530,794 | 1,507,596 | 14,084,143 | 6,242,456 | 1,251,077 | 894,350 | 635,005 | 21.42% | 2,475 |
| | Testis#3 | HK05 | 8.2 | 16,135,021 | 15,352,462 | 1,160,350 | 2,319,774 | 11,872,338 | 3,769,704 | 617,744 | 614,821 | 517,860 | 19.98% | 1,536 |
| | Testis#4 | HK06 | 8.5 | 17,486,533 | 16,778,665 | 916,317 | 2,468,053 | 13,394,295 | 5,533,221 | 922,286 | 687,128 | 513,000 | 19.23% | 1,628 |
